# An autoactive *NB-LRR* gene causes *Rht13* dwarfism in wheat

**DOI:** 10.1101/2022.05.28.493833

**Authors:** Philippa Borrill, Rohit Mago, Tianyuan Xu, Brett Ford, Simon J Williams, Adinda Derkx, William D Bovill, Jessica Hyles, Dhara Bhatt, Xiaodi Xia, Colleen MacMillan, Rosemary White, Wolfram Buss, István Molnár, Sean Walkowiak, Odd-Arne Olsen, Jaroslav Doležel, Curtis J Pozniak, Wolfgang Spielmeyer

## Abstract

Semidwarfing genes have greatly increased wheat yields globally, yet the widely used gibberellin (GA) insensitive genes *Rht-B1b* and *Rht-D1b* have disadvantages for seedling emergence. Use of the GA sensitive semidwarfing gene *Rht13* avoids this pleiotropic effect. Here we show that *Rht13* encodes a *nucleotide-binding site/leucine-rich repeat (NB-LRR)* gene. A point mutation in the semidwarf *Rht-B13b* allele autoactivates the *NB-LRR* gene and causes a height reduction comparable to *Rht-B1b* and *Rht-D1b* in diverse genetic backgrounds. The autoactive *Rht-B13b* allele leads to transcriptional upregulation of *pathogenesis-related* genes including *class III peroxidases* associated with cell wall remodelling. *Rht13* represents a new class of reduced height (*Rht*) gene, unlike other *Rht* genes which encode components of the GA signalling or metabolic pathways. This discovery opens new avenues to use autoactive *NB-LRR* genes as semidwarfing genes in a range of crop species, and to apply *Rht13* in wheat breeding programmes using a perfect genetic marker.

## Introduction

Dwarfing or reduced height genes have been associated with large increases in the yield of cereals since they were introduced during the Green Revolution (Hedden, 2003). Most current wheat cultivars carry *Rht-B1b* or *Rht-D1b* which encode negative regulators of gibberellin (GA) signalling (Peng *et al*., 1999), resulting in GA insensitivity and reduced height. These GA insensitive alleles confer benefits to yield by optimising resource partitioning to the grain and reduced lodging (Thomas, 2017). However they have pleiotropic effects on growth including reductions in coleoptile length and seedling leaf area (Allan, 1980) and impact resistance to diseases such as fusarium head blight (Srinivasachary *et al*., 2009). The use of alternative dwarfing genes that do not disrupt GA signalling, and which can reduce final plant height without adverse effects on seedling growth, will be particularly relevant in water limited environments (Richards *et al*., 2010).

Several alternative dwarfing loci have been discovered (McIntosh *et al*., 2020) which are GA sensitive and could therefore overcome the limitations of *Rht-B1b* and *Rht-D1b* on early growth. Recently, the causal genes for some of these alternative dwarfing loci have been identified, revealing their functions in the GA metabolic pathway. The first of these to be identified was *Rht18*, which is on chromosome 6A and causes an increased expression of a *GA 2-oxidase* gene (*GA2oxA9*) resulting in the removal of GA_12_ precursors from the GA biosynthesis pathway, a reduction of bioactive GA_1_ and reduced plant height (Ford *et al*., 2018). Map position, allelism tests and increased expression of the same *GA 2-oxidase* gene in *Rht14* lines suggested that *Rht14* and *Rht18* are allelic (Haque *et al*., 2011; Tang, 2016; Ford *et al*., 2018). Increased expression of related *GA 2-oxidase* genes was also found to be responsible for other alternative dwarfing alleles such as *Rht12* (*GA2oxA13* on chromosome 5A) (Sun *et al*., 2019; Buss *et al*., 2020) and *Rht24* (*GA2oxA9* on chromosome 6A, not allelic with *Rht18*) (Tian *et al*., 2022). These alternative dwarfing genes appear to operate through a shared mechanism, i.e. reduction of the flux through the GA biosynthetic pathway and subsequently lower GA content. In addition to *GA 2-oxidase* genes on chromosome 5A and 6A, other *GA 2-oxidase* genes were identified in the wheat genome (Pearce *et al*., 2015), suggesting that other dwarfing genes at different positions may also cause increased expression of other members of the *GA 2-oxidase* family.

*Rht13* is another promising alternative dwarfing gene that reduces final plant height without affecting seedling growth (Ellis *et al*., 2004; Rebetzke *et al*., 2011). The dwarfing allele *Rht-B13b* produced a strong height reduction between 17 % and 34 % compared to *Rht-B13a*, which is comparable to reductions typical of *Rht-B1b* and *Rht-D1b*, depending on the genetic background and growing conditions (Rebetzke *et al*., 2011; Rebetzke *et al*., 2012; Wang, Y *et al*., 2014; Wang *et al*., 2015; Divashuk *et al*., 2020). Genetic mapping located *Rht13* on the long arm of chromosome 7B (Ellis *et al*., 2005) but the underlying gene has not yet been identified. Here we describe the causal gene that encodes an autoactive allele of a *nucleotide-binding site/leucine-rich repeat (NB-LRR)* gene at the *Rht13* locus on chromosome 7BL. Autoactivation of *Rht13* leads to upregulation of pathogenesis-related (*PR*) genes, including class III peroxidases, which may catalyse the cross-linking of cell wall compounds to limit cell elongation and hence reduce height.

## Methods

### Introduction of Rht13 dwarf allele into different genetic backgrounds

The *Rht13* dwarfing gene was originally generated by C.F. Konzak at Washington State University in the 1980s by treating the Argentinian wheat Magnif 41 (PI344466) with N-methyl-N’ nitrosourea and selecting a semidwarf line Magnif 41M1 (Konzak, 1982). Seed of Magnif 41 (AUS17236) and Magnif 41 M1 (AUS17520) was obtained from Winter Cereal Collection, Tamworth, Australia. Magnif M41 (subsequently called Magnif) and Magnif M41 M1 (subsequently called Magnif M) plants were grown in 20 cm pots containing compost in the glasshouse maintained under 16 hr light, 23°C day and 16°C night. Internode lengths were measured at maturity for five plants per genotype.

Magnif M was backcrossed into three adapted Australian cultivars (EGA Gregory, Espada and Magenta) and BC_1_F_3_ plants were screened to generate homozygous BC_1_F_4_ lines carrying either a dwarfing allele (*Rht-B13b, Rht-B1b* or *Rht-D1b*) or no dwarfing alleles (wild type alleles: *Rht-B13a, Rht-B1a* and *Rht-D1a*). Seed of 1-4 independent F_4_ sister lines was increased from each genotype and background to generate BC_1_F_5_ seed for planting in rows in a field with bird-proof netting (birdcage), Canberra in 2014. Three rows (20 plants/row) of each genotype/background combination were planted and 5-20 plants were measured for final height and peduncle length at maturity. Rows of Magnif and Magnif M were also included in the birdcage experiment.

### Genetic mapping of Rht13

A mapping population was generated from a cross between a homozygous short line carrying *Rht13* (ML45-S) and homozygous tall line (ML80-T); these lines were selected progeny from a cross between Magnif M and a tall Russian experimental line LAN. Approximately 2,400 F_2_ gametes from ML45-S x ML80-T population were screened by capillary electrophoresis using simple sequence repeat (SSR) markers *gwm577* (*wms577*) and *wmc276* that were previously shown to flank the locus (Ellis *et al*., 2005). By extracting DNA from half seeds, recombinant embryos were selected for planting and events were fixed in the F_3_ generation before homozygous lines were phenotyped for height in the glasshouse.

To identify additional markers within the genetic interval, parental lines were screened first with the 9K SNP array (Cavanagh *et al*., 2013) and then with the 90K array (Wang, S *et al*., 2014) using the genotyping platform at Agriculture Victoria Research, Bundoora, Victoria. Additional markers were validated in the recombinants using kompetitive allelic specific PCR (KASP) assays derived from array markers and additional PCR-based markers (Supplementary Tables 1 and 2). Fine mapping of *Rht13* benefited from early access in 2013 to the physical map of 7B from Chinese Spring that was based on the assembly of bacterial artificial chromosomes (BACs) from flow sorted chromosomal DNA and coordinated by University of Life Sciences in Norway and the International Wheat Genome Sequencing Consortium (IWGSC *et al*., 2018). We utilised BAC contigs and sequences to generate new markers in the target interval (Supplementary Tables 1 and 2). Two markers *7J15*.*144I10_2_2* and *127M17*.*134P08_3* that were derived from BAC sequences that flanked the locus were used to delineate the target region in the whole genome assembly of CDC Stanley (see Methods: Chrom-seq).

A second population from the cross Magnif x Magnif M was generated where the *Rht13* mutation segregated in a homogenous background. Thirty three F_3_:F_4_ lines were tested to ensure homozygosity at *Rht13* before four short lines and two tall lines were selected for Chrom-seq and RNA-seq experiments, with an additional two tall lines included in the RNA-seq experiments to bring the total to four tall lines (see Methods: RNA-seq analysis for candidate gene identification). The KASP marker which was developed from the functional SNP at *Rht13* co-segregated with height in 33 homozygous F_4_ lines (see Methods: Validation of *Rht13* in Cadenza mutant).

### Chromosome-sequencing

To purify and sequence chromosome 7B, we selected four short and two tall progeny derived from the Magnif x Magnif M cross which had been tested to ensure homozygosity at *Rht13*. Briefly, suspensions of mitotic metaphase chromosomes were prepared from synchronized root tip meristem cells following Vrána *et al*. (2000) and Kubaláková *et al*. (2005). Prior to the flow cytometric analysis, chromosomes were labelled by fluorescence *in situ* hybridization in suspension (FISHIS) using 5’-FITC-GAA_7_-FITC-3’ oligonucleotide probe according to Giorgi *et al*. (2013) and stained by DAPI (4’,6-diamidino 2-phenylindole) at 2 µg/mL. Chromosome analysis and sorting was done using FACSAria II SORP flow cytometer and sorter (Becton Dickinson Immunocytometry Systems, San José, USA). DAPI vs. FITC dot plots were acquired for each sample (Supplementary Figure 1) and chromosomes were sorted at rates of 1500 -2000 particles per second. 50,000 - 70,000 copies of 7B chromosomes were sorted from each genotype into PCR tubes containing 40 μL sterile deionized water. The sorted chromosome samples were treated with proteinase K, chromosomal DNA was purified and amplified to 5.4 - 7.9 μg by multiple displacement amplification (Supplementary Table 3) using an Illustra GenomiPhi V2 DNA Amplification Kit (GE Healthcare, Chalfont St. Giles, United Kingdom) as described by Šimková *et al*. (2008). Chromosome content of the sorted fractions was estimated by microscopic analysis of 1500 - 2000 chromosomes sorted onto a microscopic slide. After air-drying, chromosomes were labelled by FISH with probes for pSc119.2 and Afa family repeats (Molnár *et al*., 2016) and least 100 chromosomes from each sort run were classified following the karyotype of Kubaláková *et al*. (2005).

The purified DNA from chromosome 7B was sequenced using short-read Illumina 150 bp paired end reads. The raw reads from the samples were trimmed using trimmomatic v0.32 (Bolger *et al*., 2014) (parameters: ILLUMINACLIP:TruSeq3-PE.fa:2:30:10:8:TRUE LEADING:3 TRAILING:3 SLIDINGWINDOW:4:15 MINLEN:36). Trimmed reads were subsequently mapped to the IWGSC RefSeqv1.0 Chinese Spring (IWGSC *et al*., 2018) and the CDC Stanley reference genome sequence (Walkowiak *et al*., 2020) using HISAT2 v2.1.0 (Kim *et al*., 2019) (--rg id and --rg were set per sample to enable variant calling). CDC Stanley was included in the analysis due to poor mapping of Magnif reads to the Chinese Spring reference genome in the *Rht13* mapping interval. Prior to mapping we divided the CDC Stanley pseudomolecules each into two parts to make them compatible with downstream analysis software (see Supplementary Table 4 for details). The output sam file was sorted using samtools v1.8 (Li *et al*., 2009), mate pair coordinates added using samtools fixmate, duplicates removed using samtools markdup and reads mapping to chromosome 7B part2 were selected using samtools view. We used freebayes v1.2.0 (Garrison & Marth, 2012) to call variants between the samples and the CDC Stanley reference sequence on chromosome 7B part2, with settings in freebayes only keeping variants with 2 alleles, using reads with a MAPQ>7 and a base quality >20 (--use-best-n-alleles 2 --min-mapping-quality 7 --min-base-quality 20). We compared the flanking marker sequences (obtained from the BAC sequences) to CDC Stanley using blastn in BLAST v2.9.0 (Camacho *et al*., 2009) and kept the best hit for each flanking marker (all >98% ID). This enabled the identification of the physical sequence for the *Rht13* mapping interval in the CDC Stanley genome. We filtered the freebayes output using vcftools v0.1.15 to retain variants present within the mapping interval (--from-bp 339467956 --to-bp 341325941), variants which had 2 alleles (i.e. all samples did not have the same non-ref allele, --min-alleles 2) and variants with at least 3 reads mapping (--min-meanDP3). We manually inspected the vcf file to identify homozygous variants between tall and short plants. In total we identified 13 variants within the mapping interval which were homozygous for one allele in tall lines and homozygous for a different allele in the short lines.

### Alignment between Chinese Spring and CDC Stanley chromosome 7B

Whole chromosome alignments were performed for chromosome 7B of Chinese Spring and CDC Stanley using MUMmer v4.0 (Marçais *et al*., 2018) and the nucmer command, with minimum match set to 1000. For a localized alignment of the *Rht13* region between 705 and 725 Mbp, the minimum match was set to 100. In both cases, the alignments were filtered for the best alignment, in the case of multiple alignments, and then filtered for a percent identity of 98% or greater. Dotplots were then generated using mummerplot and visualized in gnuplot v4.6.

### RNA-seq analysis for candidate gene identification

We used the same four short and two tall progeny segregating from a Magnif x Magnif M cross for RNA-seq that were used for Chrom-seq. We included an additional two tall progeny from the same population which had been tested to ensure homozygosity at *Rht13*. The basal 25% of elongating peduncles from the main stem were harvested at 50% final length and immediately frozen in liquid nitrogen. RNA was extracted using Qiagen RNeasy kit and sequenced using Illumina 150 bp paired end reads. The reads for each sample were trimmed with trimmomatic v0.32 using the same parameters as for the chrom-seq reads. The trimmed reads were then aligned to the CDC Stanley pseudomolecules (with each chromosome divided into two parts) using HISAT2 v2.1.0 with the option –dta to facilitate downstream transcript assembly using StringTie (Pertea *et al*., 2015). Transcripts were assembled using StringTie v1.3.3 for each sample individual, before merging the transcript assemblies using StringTie --merge. This produced 70,317 transcripts across all eight samples. We calculated abundance for each transcript per sample using StringTie (parameters: -e -B) and extracted the count data using the StringTie python script prepDE.py. Upon examination, a principal component analysis plot revealed that one of the Magnif samples was an outlier from the other three replicates, so this sample was excluded from further analysis. Differentially expressed genes were identified using DESeq2 1.26.0 (Love *et al*., 2014), with differentially expressed genes defined as padj <0.001. Only six transcripts were contained in the *Rht13* mapping interval and only 1 transcript was differentially expressed. We cross-referenced whether these six transcripts contained any of the 13 variants identified by chrom-seq.

### Annotation of candidate gene as NB-LRR

The candidate gene (*MSTRG*.*55039*) was annotated on the CDC Stanley reference assembly (Supplementary file 1). The longest protein sequence was identified using a three frame forward and reverse translation of the transcript using Expasy (Artimo *et al*., 2012), this protein is provided in Supplementary file 1. We searched the NCBI database using blastp for similar protein sequences, all of the top hits were NB-LRR genes, but the maximum percentage ID was only 66.5 %. We identified the position of the NB-ARC and LRR domain using the NCBI conserved domain database (Marchler-Bauer *et al*., 2017) (Supplementary file 1).

### RNA-seq analysis to understand biological role of Rht13

Using the DESeq2 results, we considered genes to be differentially expressed where padj <0.001 and expression was >2 fold up/downregulated between short and tall samples. *Pathogenesis related* (*PR*) gene sequences reported in Zhang *et al*. (2017) were downloaded from NCBI and were identified in our StringTie transcript assembly by using blastn (BLAST v2.9.0), keeping the best hit.

### Gene Ontology (GO) term enrichment

To annotate the StringTie transcript assembly with GO terms we used blastn (BLAST v2.9.0) (Camacho *et al*., 2009) to identify the best hit in the Chinese Spring RefSeqv1.1 annotation (IWGSC *et al*., 2018). For transcripts which were >95 % identical across >200 bp, the GO terms were transferred from Chinese Spring to the StringTie assembly. In total 43,685/59,228 genes were assigned a GO term using this approach. GO term enrichment analysis was carried out separately for upregulated and downregulated genes using goseq v1.38.0 (Young *et al*., 2010). The resulting GO term list were summarised using Revigo (Supek *et al*., 2011) on the medium setting (0.7) using rice (*Oryza sativa*) GO term sizes.

### Identification of class III peroxidases

We used the list of class III peroxidase genes identified by Yan *et al*. (2019) in the Chinese Spring survey sequence. We extracted coding sequence for each class III peroxidase gene and used blastn (BLAST v2.9.0) (Camacho *et al*., 2009) to identify corresponding sequences in our StringTie transcript assembly for the Magnif samples. We filtered the results to only keep hits >95 % identical with a length >400 bp. After removing duplicate transcripts, we had 242 transcripts from 219 genes. Of these 219 genes, 29 were 2-fold upregulated padj <0.001. To confirm the identity of these 29 differentially expressed genes as class III peroxidases we used RedoxiBase (Savelli *et al*., 2019) to carry out a blastx of their transcripts against the Peroxibase curated peptide database. One of the differentially expressed genes was annotated as an ascorbate peroxidase by RedoxiBase so it was excluded, while the other 28 genes were confirmed to be class III peroxidases.

### Validation of Rht13 in Cadenza mutants

We searched for the mutation identified in Magnif M in the Cadenza TILLING population (Krasileva et al., 2017). The candidate gene was not present in the Chinese Spring reference sequence (best BLAST hit TraesCS7B02G452600, 79 % identity) so we could not use the mapped mutations at PlantsEnsembl (Howe *et al*., 2020). Instead, we used www.wheat-tilling.com (Krasileva *et al*., 2017) which includes mutations called on *de novo* assembled contigs, which may not be present in Chinese Spring. The best BLAST hit to the candidate gene genomic sequence was on contig TGAC_Cadenza_U_ctg7180000823280, which had 100 % identity across 3,987 bp, including the entire CDS. We annotated the gene present on this contig and we identified Cad0453, which contained the same point mutation resulting in the identical amino acid as the Magnif M lines (Supplementary file 1). We used Polymarker (Ramirez-Gonzalez *et al*., 2015b) to develop a primer to distinguish the wild type and mutant allele using KASP genotyping (LGC Genomics). The primer sequences were: forward primer mutant (*Rht-B13b*) allele: ctgctatgggtgtgcgtctT, forward primer wild type (*Rht-B13a*) allele: ctgctatgggtgtgcgtctC, common reverse primer: cctctcacgagctgcttcaa. The standard FAM/HEX compatible tails were added at the 5’ end and the target SNP was present at the 3’ end (Ramirez-Gonzalez *et al*., 2015a).

### Phenotyping and genotyping of Cadenza0453 mutants

Twenty seeds from the M_5_ line of Cadenza0453 was grown in a growth chamber with 16 hr light, 20°C day, 16°C night. DNA was extracted following a protocol from www.wheat-training.com (Training, nd), adapted from Pallotta *et al*. (2003). KASP assays were performed as previously described (Ramirez-Gonzalez *et al*., 2015a) using the primers above. Plant height was measured once final height was reached (Zadoks stage 85). Ear, peduncle, and individual internode lengths were recorded for six homozygous wild type (*Rht-B13a*) individuals and eight homozygous mutant (*Rht-B13b*) individuals.

### Validation of Rht13 transgene in Fielder background

#### Constructs and transformation

The pVecBar-Rht13 construct contained a 6,998 bp fragment including 2,532 bp upstream and 450 bp downstream regions amplified from Magnif mutant genomic DNA using primers Rht13-NotF2 (5’ AATGCGGCCGCAATCGATAGGAGAGCTGCGTCTGTGTG 3’) and Rht13-AscR2 (5’ TGCGTACGGCGCGCCGAGAGTCGCCTTGCCAGTTC 3’) with Phusion® High-Fidelity DNA Polymerase (NEB, USA). pVecBarIII is a derivative of pWBvec8 (Wang *et al*., 1998), in which the 35S hygromycin gene was replaced by the bialaphos resistance gene (bar). The wheat cultivar Fielder was transformed using the *Agrobacterium tumefaciens* strain GV3101 (pMP90) as described in Richardson *et al*. (2014). T_0_ and T_1_ transformants were tested for the presence of transgenes by PCR using primers F698 (5’ AGGTCCTTGTGACCGAAATG 3’) and R1483 (5’ CAGTGAGCCTTTCCTGTTCC 3’).

To identify the copy number of transgenes in transgenic plants, genomic DNA from individual T_1_ segregating plants from transgenic events were used for DNA gel blot hybridisation as described in Mago *et al*. (2015). DNA was digested with HindIII and a part of the selectable marker gene ‘bar’ was used as a probe.

#### Gene expression

Expression of the transgene was done using qRT-PCR analysis. Leaf tissue was collected from individual plants of a segregating T_1_ family at stem elongation stage (Zadoks stage 33). RNA extraction was done using RNeasy kit (Qiagen) according to manufacturer’s instructions. Quantitative PCR was carried out on a Bio-Rad CFX96 Touch Real-Time PCR Detection System (Bio-Rad) using iTaq universal SYBR Green supermix (Bio-Rad) and a two-step cycling program according to the manufacturer’s instructions and as described in Moore *et al*. (2015). Minus RT controls were first tested with housekeeping gene *TaCON* (Moore *et al*., 2015) to ensure amplification of residual genomic DNA was insignificant. Primers qrht13-2F: 5’ GCAAAGGTTGAACTACTGTTCC 3’ and qrht13-2R: 5’ AACATCACAAAACGAACATGGA 3’ were used for quantification of *Rht13* transcript. The green channel was used for data acquisition. Efficiency and cycle threshold values were calculated using the LinRegPCR quantitative PCR data analysis (Ruijter *et al*., 2009), and relative expression levels were calculated using the relative expression software tool (REST) method (Pfaffl, 2001) relative to the housekeeper gene *TaCON*.

#### Phenotyping

For phenotyping of the transgenic plants, ten T_1_ progeny seeds from 4 independent T_0_ plants were sown in 13 cm pots containing compost in a glasshouse maintained under 16 hr light, 23°C day and 16°C night. Plant height was measured at maturity (∼Zadoks’ stage 70-80).

### Transient expression in tobacco

The coding sequence for the wild type (*Rht-B13a*) and mutant (*Rht-B13b*) allele of *Rht13* were synthesised (Twist Bioscience) and cloned into the Gateway binary vector pGWB12 with an N-terminal FLAG tag (Nakagawa *et al*., 2007). These were transformed into *Agrobacterium tumifaciens* (strain AGL-1) by electroporation. Transformed colonies were selected from agar plates supplemented with 50 μg/ml kanamycin and 50 μg/ml rifampicin and inoculated into liquid LB media with 50 μg/ml kanamycin and 50 μg/ml rifampicin. Cultures were incubated at 28°C in a shaking incubator for 24 hrs. Agrobacterial cells were harvested by centrifugation and resuspended in MMA solution [10 mM MES (2-[N-morpholino]ethanesulfonic acid) at pH 5.6, 10 mM MgCl_2_ and 150 μM acetosyringone] to a OD_600_ of 3. After incubation in the dark for 1 hr, the agrobacterial suspension was infiltrated into 4 to 5 week old *Nicotiana bethamiana* leaves. Photographs were taken 6 days after infiltration. The *N. benthamiana* plants were grown in M3 compost (Levington) mixed 3:1 with perlite under 12 hr light at 20°C day, 16°C night in a growth cabinet.

### Pathogenesis-related (PR) gene expression

Cadenza0453 plants that were homozygous for the *Rht13* wild type (*Rht-B13a*) or homozygous mutant (*Rht-B13b*) allele were grown as described above. Tissues were harvested at seven days after anthesis and snap frozen in liquid nitrogen. Four biological replicates were harvested for each tissue: flag leaf blade (central 3 cm), basal peduncle (bottom 3 cm, flag leaf sheath removed before snap freezing) and apical peduncle (top 3 cm of peduncle tissue just below the rachis node). RNA was extracted using the RNeasy Plant Mini Kit (Qiagen) according to the protocol from the manufacturer, using the RLT buffer. Genomic DNA was digested by RQ1 RNase-free DNAse (Promega) according to the manufacturer’s instructions. cDNA was synthesised using the AffinityScript Multiple Temperature cDNA Synthesis Kit (Agilent) with random primers according to the manufacturer’s instructions with a synthesis temperature of 55°C.

qPCR was carried out with 3-4 biological replicates with 3 technical replicates per reaction. Primers for *PR3* and *PR4* were from Zhang *et al*. (2017) and for *actin* were from Uauy *et al*. (2006). qPCR was carried out using PowerUp SYBR Green (Applied Biosystems) according to the manufacturer’s instructions with each primer at a final concentration of 0.25 µM and 0.5 µL of cDNA in a 10 µL reaction, using 384 well plates. The qPCR programme run on the QuantStudio5 (ThermoFisher) was as follows: pre-incubation at 50°C for 2 min and 95°C for 2 min; 40 amplification cycles of 95°C for 15 s, 58°C for 15 s, and 72°C for 1 min. The final melt-curve step heated to 95°C for 15 s, cooled to 60°C for 1 min and then heated to 95°C with continuous reading as the temperature increased.

All qPCR reaction melt curves were inspected to have only a single product. Crossing thresholds were calculated using the QuantStudio5 software (ThermoFisher). Expression level was calculated relative to *actin* using the Pfaffl method which accounts for primer efficiency (Pfaffl, 2001). Primer efficiencies were calculated using a serial dilution of cDNA.

### Hydrogen peroxide quantification

Hydrogen peroxide content was measured in elongating peduncles (50% final length) of Magnif and Magnif M using the protocol described in Amplex Red Hydrogen Peroxide Kit (Invitrogen). Plants were grown in a glasshouse as described in the section “Methods: Introduction of *Rht13* dwarf allele into different genetic backgrounds”. 30 mg of ground tissue was resuspended in the reaction buffer and spun down before adding it to the reaction mixture. Fluorescence was detected at 590 nm after 30 min incubation. The experiment using 3 replicates was repeated and both experiments with 6 replicates in total were used for the Student’s t-test.

### Cell length measurements

Magnif M and Magnif were grown in the glasshouse as described in the previous section, until the peduncles were fully-expanded and five individual plants of each genotype were used for analysis. One 10 cm segment was collected from the most basal part of each peduncle from the primary tiller and harvested into 70% ethanol and stored at 4°C. Before further cell length analysis, all peduncles were cleared for up to 14 days in 10% household bleach, then transferred back into 70% ethanol and stored at 4°C.

For analysis of epidermal cell lengths, a 1-cm-long segment was cut from the top of each segment of harvested, cleared peduncle, i.e. a segment between 9 and 10 cm from the base of the peduncle. Segments were transferred through 4 changes of 100% dry ethanol then dried in a Tousimis Autosamdri critical point drier and mounted on stubs for examination using a Zeiss EVO LS15 scanning electron microscope. Epidermal cell lengths were detected using the backscatter detector with 30 kV accelerating voltage in 10 Pa chamber pressure (Talbot & White, 2013). Cell lengths were measured using Zeiss Zen Blue software and analysed in MS Excel. Two distinct cell types were measured: inter-hair cells and single cells (Supplementary Figure 2).

### Analysis of physical properties of peduncles

Magnif M and Magnif were grown to maturity in a glasshouse maintained at 23°C day-time temperature, with 18°C night-time temperature, as described previously. Fully mature, dried stems were used for testing to avoid confounding effects of water content, with 11-12 independent primary stems sampled per genotype. A three-point bend test was carried out on the peduncles to determine bending rigidity and bending strength as described in Hyles *et al*. (2017).

### Peduncle histochemical analysis

Cadenza0453 plants that were homozygous wild type (*Rht-B13a*) or homozygous mutant (*Rht-B13b*) were grown under speed breeding conditions in a controlled environment cabinet: 22 hrs light, 2 hrs dark, 20°C day, 15°C night, 70% humidity. Peduncles were harvested 3-7 days after anthesis. Fresh sections were cut by hand from the peduncle using a razor blade from three regions: the apical peduncle immediately under the node to the ear, the mid-point of the peduncle half way between the ear and the flag leaf node, and the basal part of the peduncle just above the flag leaf node (the flag leaf sheath was removed). The sections treated with toluidine blue O or phloroglucinol-HCl as described in Pradhan Mitra and Loqué (2014) and imaged with bright-field illumination (magnification of 20X).

## Results

### Characterisation of Rht13 phenotype in Magnif

The *Rht13* semidwarfing gene was originally identified as an induced mutant in the Magnif background (Konzak, 1982). We carried out a detailed characterisation and found that *Rht13* caused a 30-35% height reduction in both greenhouse and field conditions (birdcage) (Figure 1). A comparison of internode lengths showed that most of the height reduction occurred in the peduncle and this effect was confirmed in field grown plants that were measured for height from early stem elongation to maturity (Figure 1c, d). Height differences were apparent after Zadoks growth stage 50, with reduced peduncle length accounting for most of the effect.

**Figure 1.**
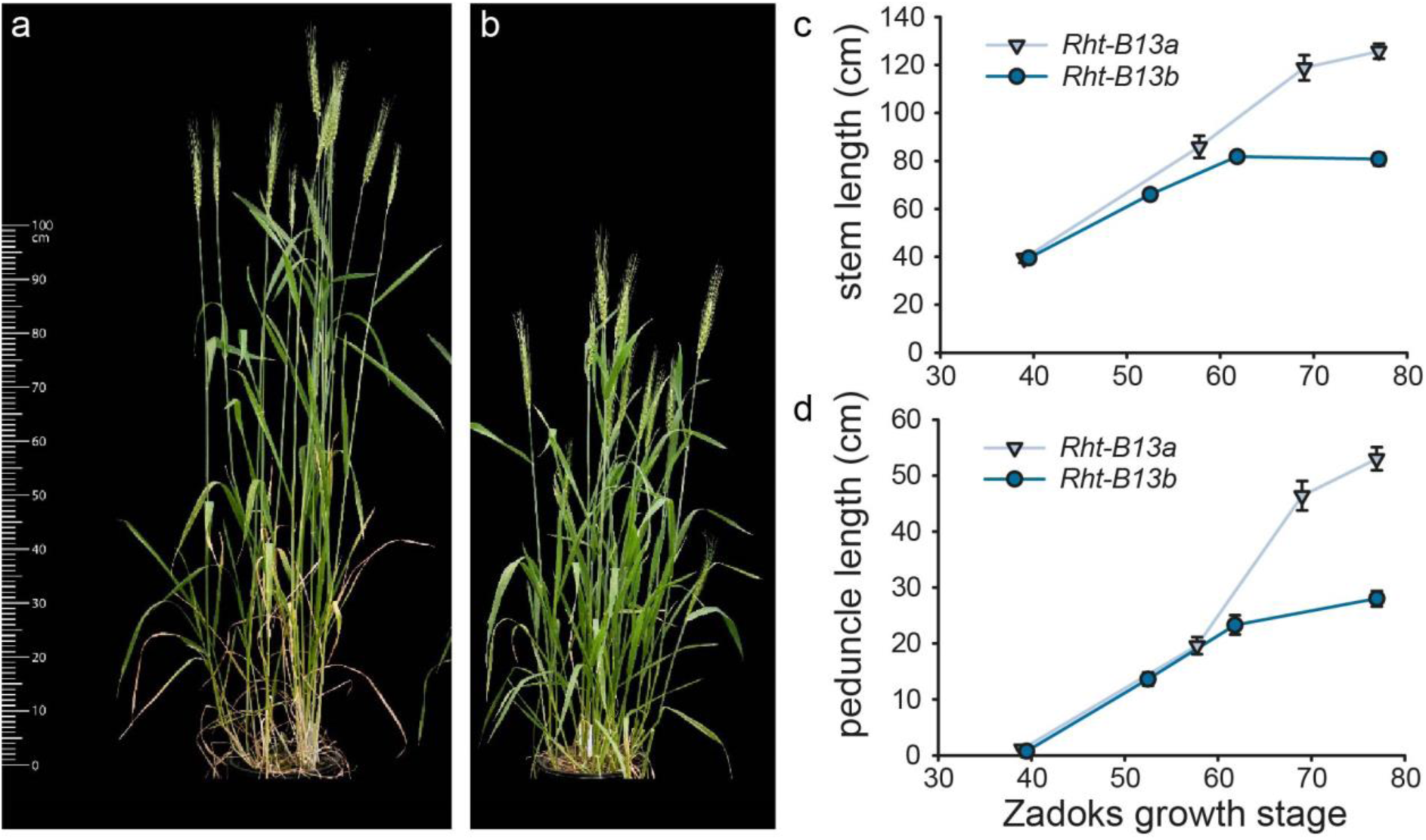
Phenotypic characteristics of Magnif (*Rht-B13a*) and Magnif M (*Rht-B13b*). a) Magnif and b) Magnif M grown under greenhouse conditions. Developmental time-course of c) stem length and d) peduncle length in wheat grown under field conditions. Data points combine measurements from 5-10 individual field grown plants. The error bars represent the standard error of the mean.

### Fine genetic mapping of Rht13 to a region on chromosome 7B

Previously, *Rht13* was mapped to the long arm of chromosome 7B and genetically linked to SSR marker *gwm577* (Ellis *et al*., 2005). An F_2_ population from a cross between parental lines ML45-S carrying *Rht13* and tall line ML80-T was developed for fine mapping.\ Approximately 2,400 F_2_ gametes were screened with SSR markers *gwm577* and *wmc276* that were previously shown to flank the locus. The screen identified 21 recombinants that corresponded to less than 1 cM of genetic distance between flanking markers (Figure 2a, Supplementary Table 2). Additional DNA markers were added to the genetic interval after parental lines were screened with the 9K and 90K wheat SNP arrays (Cavanagh *et al*., 2013; Wang, S *et al*., 2014). In addition, the project was given early access in 2013 to the emerging physical map of chromosome 7B, which was part of the international initiative to generate maps of individual Chinese Spring chromosomes led by the IWGSC and Norwegian University of Life Sciences. Several BAC clones were assigned to the region and markers that were developed from BAC sequences were added to the interval (Supplementary Table 2). BAC sequence-derived markers *7J15*.*144I10_2_2* and *127M17*.*134P08_3* flanked the *Rht13* locus on the proximal and the distal side, respectively, and defined a genetic interval of approx. 0.1 cM (Figure 2).

**Figure 2.**
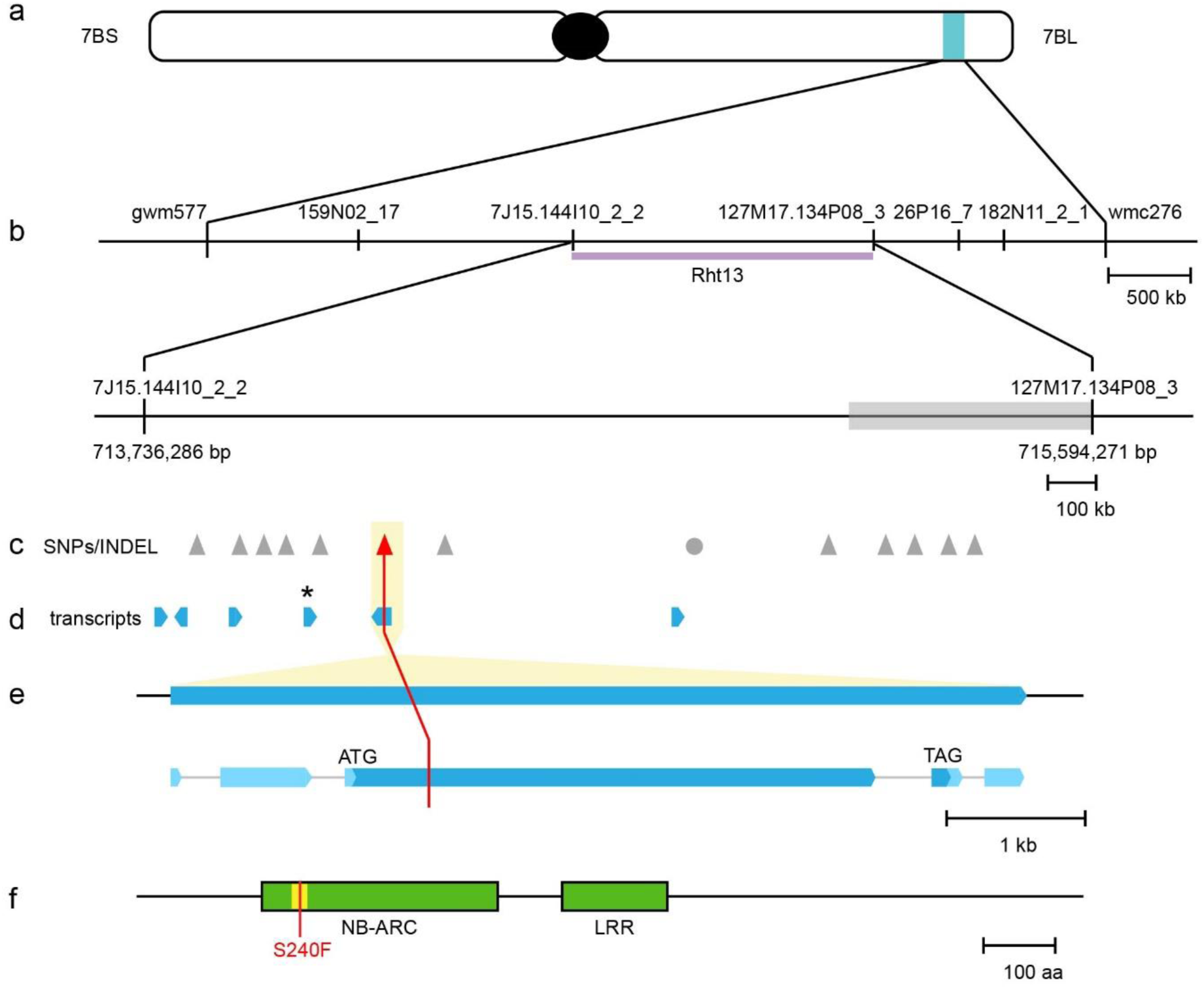
Mapping of the NB-LRR gene *Rht13*. a) *Rht13* is located on the long arm of chromosome 7B. b) Physical mapping interval in CDC Stanley with genetic markers (SSR and BAC derived). The distal region (grey box) contained more SNPs between all samples and the reference sequence. c) SNPs (triangles) and INDEL (circle) between tall and short progeny from a Magnif x Magnif M cross, identified by chrom-seq. Red triangle indicates amino acid change inducing SNP. d) Transcripts identified by RNA-seq of progeny from a Magnif x Magnif M cross. The asterisk indicates a significantly differentially expressed transcript between tall and short progeny. e) Intron-exon structure of gene encoded by *Rht13*. Exons are represented by boxes, with untranslated regions in pale blue and coding regions in darker blue, introns are represented by thin grey lines. f) The gene encodes a 1,272 amino acid protein containing an NB-ARC and LRR domain and is annotated as MSTRG.55039 (Supplementary File 1). Magnif M has a mutation (S240F) in the RNBS-A motif (yellow).

### Next generation sequencing approaches revealed a single amino acid change between expressed genes in the region on chromosome 7B

The *Rht13* region defined by flanking markers *7J15*.*144I10_2_2* and *127M17*.*134P08_3* corresponded to a 1.93 Mb interval on chromosome 7B in Chinese Spring RefSeqv1.0. To identify candidate SNPs in the interval, we generated an additional population from a Magnif x Magnif M cross and selected four short and two tall F_3_:F_4_ lines that were homozygous at *Rht13*. For each of these lines, we isolated chromosome 7B by flow sorting and then sequenced the chromosome using Illumina short-reads. We attempted to identify SNPs within the mapping interval by mapping this chrom-seq data to the RefSeqv1.0 genome sequence (IWGSC *et al*., 2018) but we found that over half of the 1.93 Mb interval had few reads mapping (1.07/1.93 Mb), which suggested haplotype divergence between Chinese Spring and Magnif. We then examined the alignment of chromosome 7B between Chinese Spring and several cultivars whose genome sequences were available from the 10+ Wheat Genomes Project (Walkowiak *et al*., 2020). We found that CDC Stanley had significant haplotype divergence from Chinese Spring in the *Rht13* interval on chromosome 7B (Supplementary Figure 3); therefore we tested whether CDC Stanley would be a more appropriate reference sequence. Using CDC Stanley as the reference, the flanking markers spanned 1.86 Mb on chromosome 7B (Figure 2b). Within this interval, a 0.49 Mb region had more SNPs between all samples and the reference sequence, suggesting some divergence between CDC Stanley and Magnif.

We identified 12 SNPs and 1 INDEL between the tall and short fixed lines in the mapping interval (Figure 2c). To identify potential causal genes for *Rht13*, we carried out RNA-seq on developing peduncle tissues from four fixed short and four fixed tall F_3_:F_4_ lines from the same Magnif x Magnif M population that was used for chrom-seq. We found that one transcript within the interval was more highly expressed in Magnif M than Magnif samples (2.5-fold change, padj <0.001; indicated by * in Figure 2d). However, this transcript did not translate to a protein longer than 76 amino acids in any frame, suggesting that pseudogenisation might have occurred. Since there were no obvious changes in expression levels of genes within the interval, except the putative pseudogene, we examined whether the SNPs detected by chrom-seq were contained within any of the transcripts. We found that only 1 SNP (G to A at chr7B:714,391,008) was located within a transcript (Figure 2e) and this SNP was predicted to cause an amino acid change within the conserved RNBS-A motif of the predicted protein sequence (Figure 2f). A KASP marker developed for the SNP co-segregated with the height phenotype in the Magnif x Magnif M population (Supplementary Figure 4).

### The amino acid change S240F reduces plant height

The expressed transcript with an amino acid change was predicted to encode a nucleotide-binding site/leucine-rich repeat (NB-LRR) protein (Figure 2f). The mutation was predicted to cause an amino acid substitution of the serine at position 240 to phenylalanine (S240F) in the RNBS-A motif (Meyers *et al*., 2003). To test whether this amino acid change caused the reduced height phenotype observed in Magnif M, we searched the Cadenza TILLING population for mutations within closely related genes (Krasileva *et al*., 2017). Line Cadenza0453 was identified as carrying a gene that was 100% identical at the nucleotide level to the mutant NB-LRR gene at the *Rht13* locus, resulting in the same amino acid change (S240F) as found in Magnif M. The KASP marker developed for the mutation segregated within progeny derived from Cadenza0453. Homozygous mutant plants (*Rht-B13b*) were on average 16.7 cm shorter than homozygous wild type plants (*Rht-B13a*) at maturity in the Cad0453 background (Figure 3a, b; p<0.001, Student’s t-test). This difference in height was reflected in shorter peduncle and internode lengths, except for the first internode (Supplementary Table 5).

**Figure 3.**
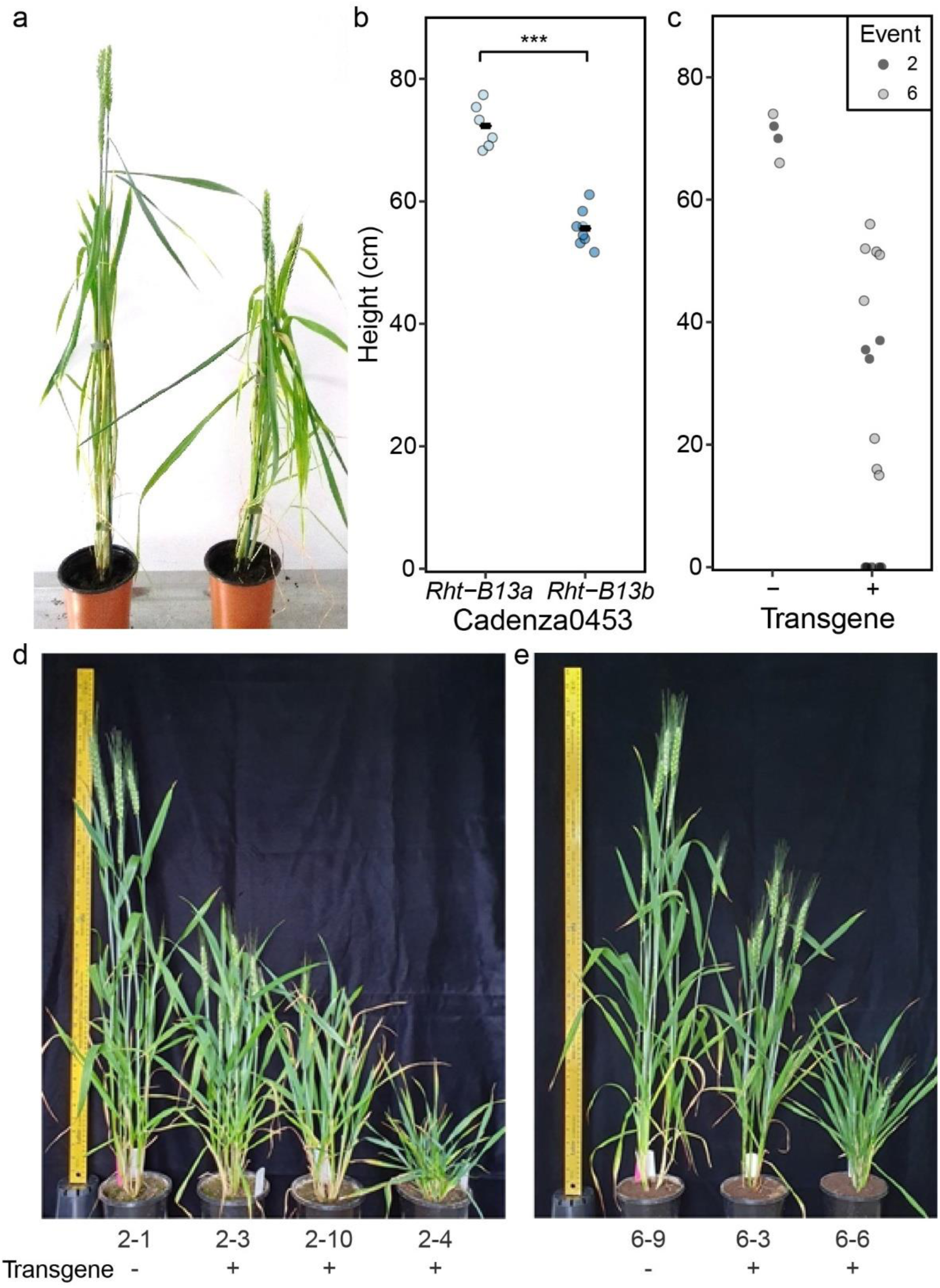
Validation that the S240F mutation in *Rht-B13b* causes a reduction in height. a) Cadenza0453 segregates for plants homozygous for the wild type allele *Rht-B13a* (left) and mutant allele *Rht-B13b* (right) and b) Cadenza0453 height quantification, the black bars represent the mean, *** p<0.001, Student’s t-test. c) Height of T_1_ progeny of two transgenic events (family 2 and 6) in Fielder background transformed with *Rht-B13b* allele, stunted plants are represented by points immediately above the x-axis (details in Supplementary Table 6). d) and e) show families 2 and 6 respectively. Null segregants (-) are on the left of each image.

To confirm that the amino acid change caused the reduction in height, we transformed the mutant allele from Magnif M (*Rht-B13b*) into Fielder (Figure 3c-e). We found that expression of the transgene caused a strong reduction in height, compared to null segregants (Figure 3c-e and Supplementary Table 6), although there was variation in the degree of dwarfism, which did not relate to the copy number or expression levels (Supplementary Figure 5 and Supplementary Table 6).

### Characterisation of the Rht13 reduced height phenotype in different genetic backgrounds

To assess the potential for use of *Rht-B13b* in breeding programmes, we generated sister lines for *Rht13* in three Australian elite backgrounds, alongside *Rht-B1b* (in EGA Gregory) *or Rht-D1b* (in Espada and Magenta) dwarfing alleles for comparison. We found that *Rht-B13b* stems elongated earlier than *Rht-B1b* or *Rht-D1b* stems, but final lengths were shorter than the tall sister lines due to an earlier arrest in growth (Figure 4a-c). This lower final length is largely due to the peduncle being shorter in *Rht-B13b* than in *Rht-B1b* or *Rht-D1b* plants (Figure 4d-f). No differences in spike length were observed. We found some differences in the effect between cultivars. In Magenta, *Rht13* is a stronger dwarfing gene than *Rht-D1b* (shorter peduncle, no difference in lower internodes; Figure 4c and f), whilst in Espada and EGA Gregory the effect of *Rht-B13b* on height is comparable to *Rht-D1b* and *Rht-B1b* (Figure 4a, b, d and e). Comparing *Rht-B13b* to tall plants lacking conventional dwarfing genes, the reductions in heights are larger in Magenta and EGA Gregory than Espada.Taken together, our results (Figures 1, 3 and 4) show that *Rht-B13b* is effective at reducing height in a range of genetic backgrounds including lines from the UK (Cadenza), Australia (Espada, EGA Gregory and Magenta), Argentina (Magnif) and the US (Fielder).

**Figure 4.**
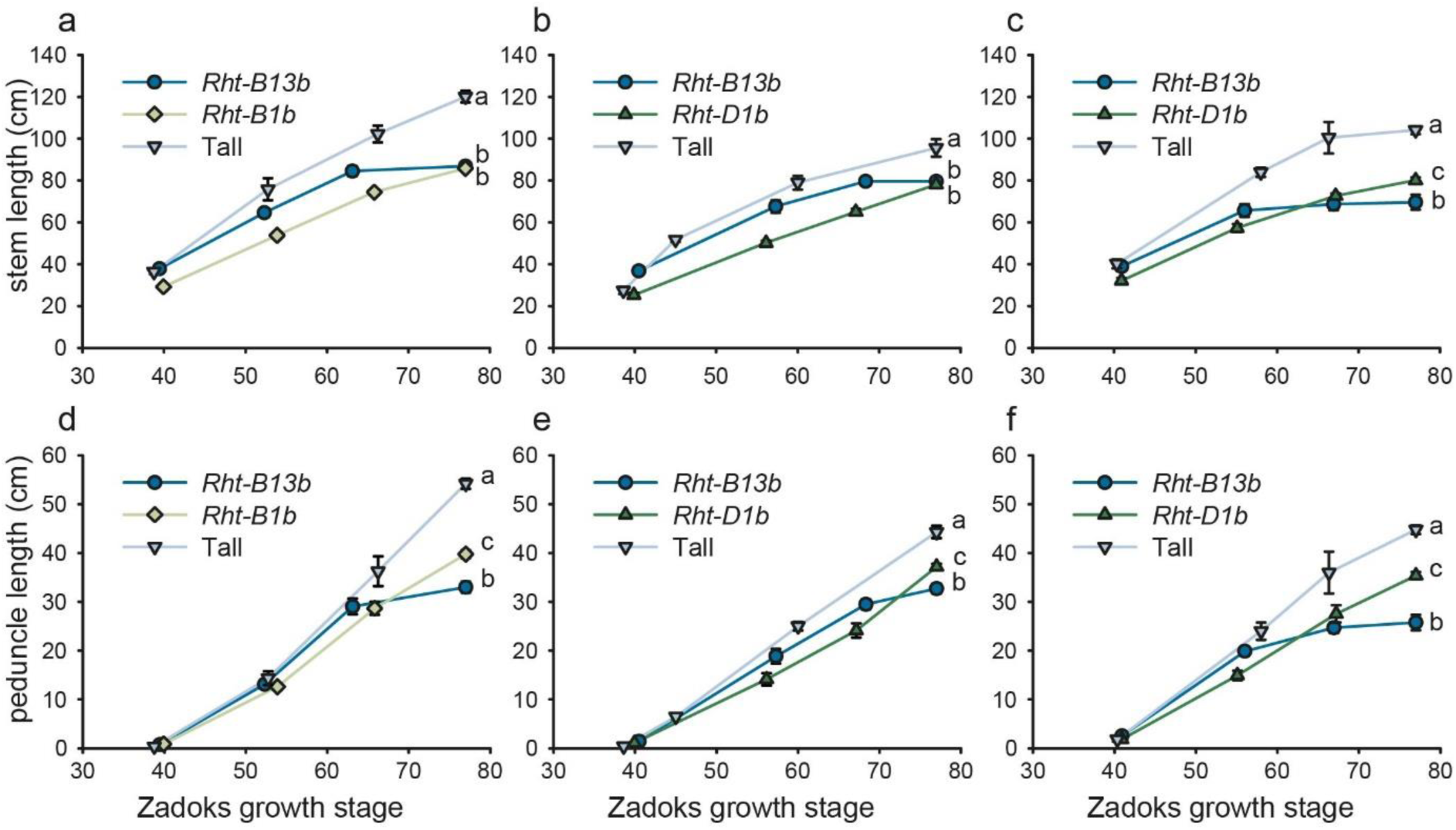
Effect of *Rht-B13b* and conventional dwarfing alleles *Rht-B1b* and *Rht-D1b* on stem and peduncle length in different wheat backgrounds in the field. a-c) stem length, d-f) peduncle length, a and d) EGA Gregory, b and e) Espada and c and f) Magenta. Letters indicate significant differences at maturity determined by a one-way ANOVA followed by Tukey post-hoc test (p < 0.05). Data points combine measurements from 5-20 individual field grown plants. The error bars represent the standard error of the mean.

### Rht-B13b is autoactive and causes a cell death response in Nicotiana benthamiana

The mutation causing the reduction in height (S240F; see figure 2f above) occurred in the RNBS-A domain of the NB-LRR protein at the same position as a mutation observed in the tomato (*Lycopersicon esculentum*) NB-LRR protein I-2 (Figure 5a). In I-2 the mutation converting the serine (S) residue to a phenylalanine (F) caused autoactivation of the protein (Tameling *et al*., 2006). Therefore, we hypothesised that the S240F mutation in *Rht13* would also result in autoactivation of the NB-LRR protein, upregulating defense responses and reducing plant growth. We first tested this through heterologous expression of the wild type (*Rht-B13a*) and mutant *Rht13* gene (*Rht-B13b*) in tobacco leaves. We found that the *Rht-B13b* allele induced more cell death 5 days post inoculation than the *Rht-B13a* allele (Figure 5b), which is a typical defense response to pathogen invasion.

**Figure 5.**
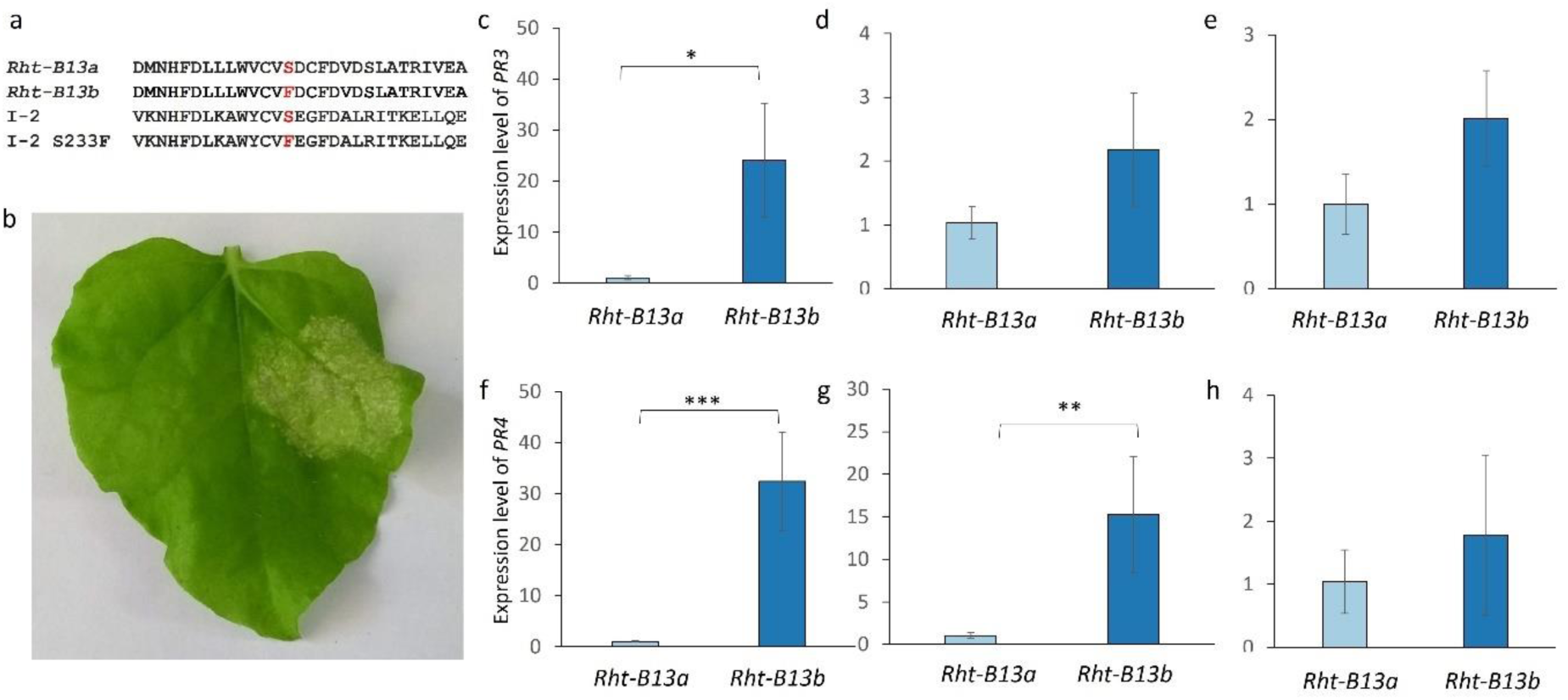
*Rht-B13b* induces defense gene responses in *Nicotiana benthamiana* and wheat. a) Alignment of the RNBS-A motif from Rht-B13a and Rht-B13b protein with the tomato I-2 protein and the I-2 mutant (S233F) that induces auto-activation. b) Infiltration of *Rht-B13b* into *N. benthamiana* induces significantly more cell death (right side of leaf) than *Rht-B13a* (left side of leaf, no cell death observed). The experiment was repeated twice, on six plants each time, a representative result is shown 6 days post inoculation. Expression of *PR* genes in wheat basal peduncle (c,f), apical peduncle (d,g) and flag leaf blade (e,h). Expression measured for *PR3* (c-e) and *PR4* (f-h) and normalised to *actin*. For each graph the expression level is normalised to be 1 in *Rht-B13a*, error bars represent the standard error (n=3-4). Significant differences were calculated using a t-test on log transformed values, * p<0.05, ** p<0.01, *** p<0.001.

Cell death responses associated with *Rht-B13b* were not observed in any of the wheat backgrounds. It is possible that autoactivation of *Rht13* in wheat might nevertheless enhance defense responses leading to a reduction in growth, without leading to cell death. We found that the expression level of p*athogenesis-related* genes (*PR* genes) *PR3* and *PR4* were >20 fold upregulated in the basal peduncle in the *Rht-B13b* mutant compared to the *Rht-B13a* wild type sibling Cadenza0453 (Figure 5c, f), suggesting that autoactivation of defense responses occurred in the *Rht13* mutant plants in rapidly expanding tissue. *PR4* was 15-fold upregulated in the apical peduncle but no significant difference was observed in *PR3* expression (Figure 5d, g). No differences were observed in *PR* gene expression between *Rht-B13b* and *Rht-B13a* in the flag leaf blade (Figure 5e, h).

### RNA-seq analysis reveals that class III peroxidases are upregulated by autoactive Rht13

To further explore the pathways through which *Rht13* reduces height we used the same RNA-seq data from peduncle samples of fixed lines from the Magnif x Magnif M population, which was previously used to identify the causal gene (see Figure 2). We confirmed that *PR* genes were upregulated in Magnif M (*Rht-B13b*) compared to Magnif (*Rht-B13a*) (Supplementary Figure 6), similar to observations in Cadenza (Figure 5c-h). The fold changes observed were higher in the RNA-seq data (Supplementary Figure 6) than the qPCR data (Figure 5c-h); however, *PR4* upregulation was only borderline significant (p=0.05). The upregulation of *PR* genes was consistent with upregulation of defense response associated genes in the Magnif M plants compared to Magnif, identified by GO term enrichment (Supplementary Figure 7). Overall, we found that more genes were upregulated (1,560 genes) than downregulated (726 genes) in Magnif M compared to Magnif (>2 fold, padj <0.001). Upregulated genes were enriched for GO terms including defense responses, cell wall organization, regulation of hydrogen peroxide metabolic processes and salicylic acid biosynthetic processes. We did not detect any enrichment for genes related to GA signalling or biosynthesis. Downregulated genes were associated with flavonoid biosynthetic processes, responses to cytokinin and photosynthesis (Supplementary Figure 7).

We further hypothesised that the autoactivation of defense responses in the mutant line will cause the production of reactive oxygen species, which can promote cross-linking and cell wall stiffening leading to less growth (Schopfer, 1996; Schmidt *et al*., 2016). To investigate this, we examined the expression of class III peroxidases that can use hydrogen peroxide in cross-linking reactions during cell wall organization and pathogen defense (Smirnoff & Arnaud, 2019). We identified 218 class III peroxidases that were expressed in Magnif or Magnif M peduncle samples. Of these, 28 were significantly upregulated in Magnif M (*Rht-B13b*) compared to Magnif (*Rht-B13a*) in the peduncle (padj<0.001, > 2-fold, Figure 6a), which is a significantly greater proportion than would be expected for a set of 218 random genes (12.8 % vs 2.6 %, Chi-squared test, p<0.001). Furthermore, many of the class III peroxidase genes were very strongly upregulated (11/28 are upregulated >10 fold).

**Figure 6.**
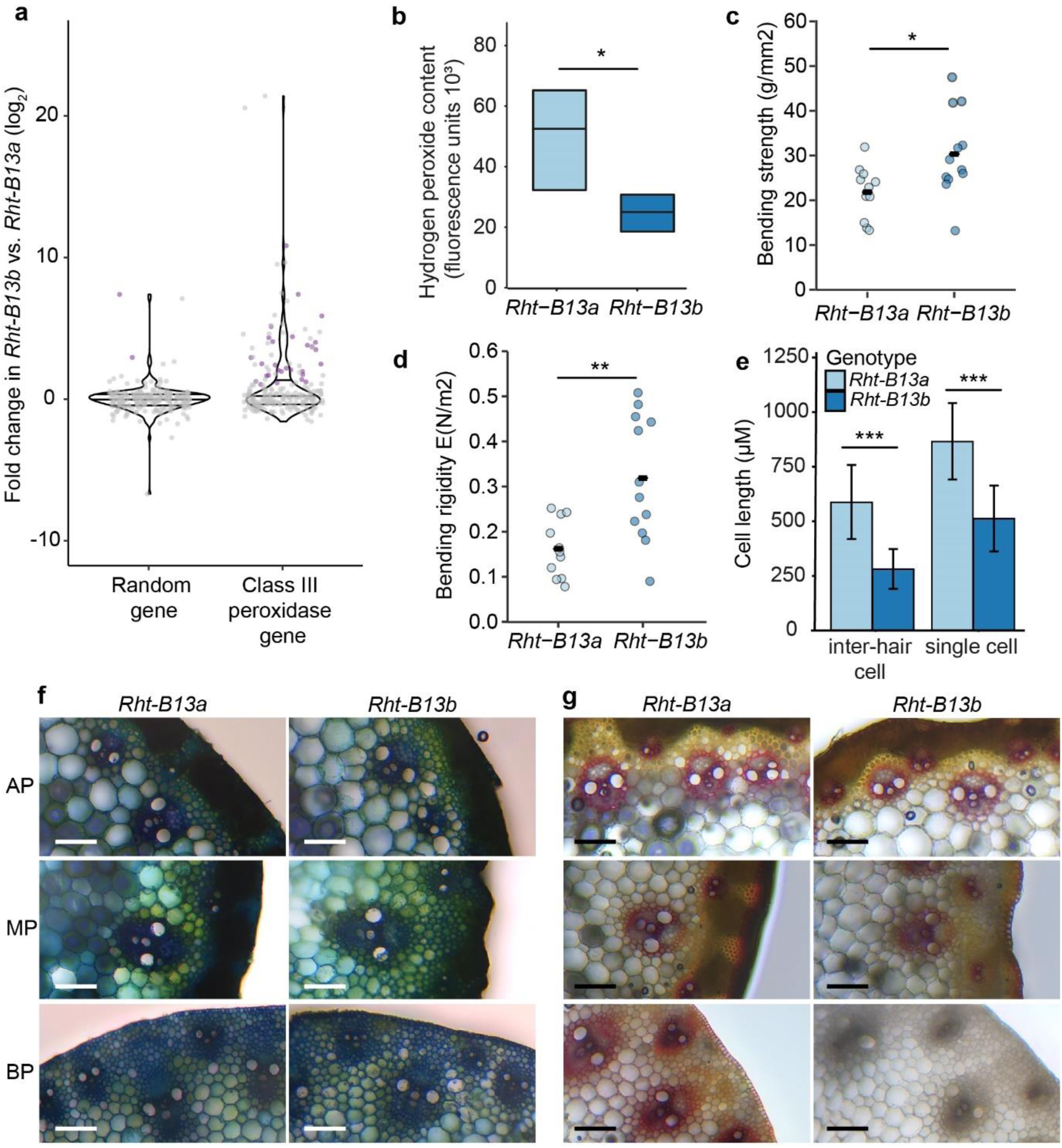
Changes in class III peroxidase gene expression, hydrogen peroxide content, mechanical and cell properties in mutant (*Rht-B13b*) compared to wild type (*Rht-B13a*) peduncles. a-e) are in a Magnif background, f) and g) are in a Cadenza background. a) Fold change in expression of 218 class III peroxidase genes compared to an equivalent number of randomly selected genes. Purple dots represent genes differentially expressed at padj<0.001 with a fold change >2, grey dots are not differentially expressed, lines across the violin plot represent quartile 1, the median and quartile 3. b) Hydrogen peroxide content in elongating peduncles. Significant differences determined by Student’s t-test, n=6. Peduncle bending strength c) and bending rigidity d) were determined using a 3-point bend test, significant differences were determined using Student’s t-tests, n=11-12. e) Epidermal cell lengths in inter-hair and single cells, significant differences determined by ANOVA, n=62-190 individual cells. f) and g) transverse sections imaged with bright-field illumination (magnification 20X) from the apical peduncle (AP) 1 cm below the ear, the peduncle mid-point (MP) and the basal peduncle (BP) 1 cm above the node. f) is stained with toluidine blue O and g) with phloroglucinol-HCl. One representative image from 5 independent biological replicates is shown. Asterisks indicate statistical differences between genotypes: * p<0.05, ** p<0.01, *** p<0.001.

We found that Magnif M (*Rht-B13b*) peduncles had lower hydrogen peroxide content than Magnif (*Rht-B13a*) (Figure 6b, p<0.05, Student’s t-test), consistent with upregulation of class III peroxidases in the mutant (Figure 6a). To test whether these gene expression and metabolite changes influence cell wall mechanical properties, we used a 3-point bend test to measure peduncle strength and ridigity. We found that the Magnif M (*Rht-B13b*) peduncles were stronger and more rigid than Magnif (*Rht-B13a*) peduncles (Figure 6c, d, p = 0.02 and p = 0.003 respectively, Student’s t-test). The Magnif M (*Rht-B13b*) peduncles had shorter cell lengths in their epidermis, with cell lengths of approximately 2/3 of wild type, suggesting a lower level of cell expansion (Figure 6e). To investigate whether these mechanical changes are mediated by changes to lignification, we examined cross sections of the peduncle taken from the apical part of the peduncle immediately under the ear, the mid-point of the peduncle, and the basal part of the peduncle just above the node. Using toluidine blue we did not observe any obvious morphological changes (Figure 6f) and no significant differences in lignification were observed between Magnif and Magnif M in the apical or middle peduncle (Figure 6g). However, the basal sections of Magnif M (*Rht-B13b*) peduncles had much lower staining of lignin in and around vascular bundles than the Magnif (*Rht-B13a*) (Figure 6g).

## Discussion

### Novel mechanism for a wheat Rht gene

A striking difference to other reported *Rht* genes in wheat is that *Rht13* is not directly involved in GA signalling or metabolism, as is the case for conventional dwarfing genes *Rht-B1b* and *Rht-D1b* (Peng *et al*., 1999) and the cloned alternative dwarfing genes *Rht12* (Buss *et al*., 2020), *Rht18* (Ford *et al*., 2018) and *Rht24* (Tian *et al*., 2022). Instead, *Rht13* is a *NB-LRR* gene with a point mutation that induces autoactivation. The amino acid change in *Rht13* is the same mutation as previously characterised in the tomato protein I-2 which impeded ATP hydrolysis and promoted an ATP-bound active form of the protein (Tameling *et al*., 2006). Due to the high conservation between the RNBS-A motif between I-2 and Rht13, we hypothesise that the mutation in *Rht13* has the same biochemical function to impede ATP hydrolysis, consistent with the hypersensitive response (HR) we observed upon expressing *Rht-B13b* in *N. benthamiana* leaves.

Autoactive *NB-LRR* genes have been reported to reduce growth in several plant species (Yang & Hua, 2004; Chintamanani *et al*., 2010; Roberts *et al*., 2013), including causing reduced internode length in flax (Howles *et al*., 2005). However, these autoactive NB-LRRs are often associated with negative pleiotropic effects including a spontaneous HR resulting in necrotic lesions. We did not observe any spontaneous HR or necrosis in any of the wheat genetic backgrounds tested. This contrasts with known autoactive *NB-LRR* genes in cereals that reduce height, such as *Rp1-D21* in maize which induces a spontaneous HR in a range of genetic backgrounds, although to differing degrees of severity (Chintamanani *et al*., 2010). Nevertheless, *Rht-B13b* induced a HR in tobacco, which could be a result of high transient expression in tobacco, although overexpression of *Rht-B13b* in wheat did not cause a HR despite severe stunting. Instead, it is possible that tissue specific expression of *Rht13* in wheat or differences in signalling pathway thresholds between tobacco and wheat may explain these differences. This is supported by our finding that *PR* genes were only upregulated in peduncle tissues, and not in the flag leaves of Cadenza *Rht-B13b*. The upregulation of *PR* genes in *Rht-B13b* containing wheat raises the question whether *Rht-B13b* could also enhance resistance response to certain pathogens. Autoactive mutants in flax, potato and tomato were shown to gain additional specificities to strains of the same pathogen or became effective against other pathogen species (Howles *et al*., 2005; Farnham & Baulcombe, 2006; Giannakopoulou *et al*., 2015), but further research will be required to determine any association between *Rht-B13b* and enhanced disease resistance.

Amongst the *PR* genes upregulated by *Rht-B13b* are class III peroxidases which are known to act in a wide range of physiological processes, including cross-linking of cell wall components, formation of lignin and metabolism of reactive oxygen species such as hydrogen peroxide (Smirnoff & Arnaud, 2019). The upregulation of class III peroxidases is associated with a decrease in hydrogen peroxide in *Rht-B13b*, which may be due to its use in cell wall cross-linking. Increased cross-linking could explain the reduced cell lengths observed in *Rht-B13b* and the increase in peduncle strength and rigidity. Surprisingly, we did not observe an increase in lignin in *Rht-B13b* compared to wild type, suggesting that these changes in cell size and tissue strength may be mediated by cross-linking polysaccharides and extensins other than lignin. Alternatively, subtle differences in lignin content may not be detectable by histochemical staining in the middle section of the peduncle, where differences in bending strength were observed. Taken together, we present a model through which *Rht-B13b* operates (Figure 7). In this model, the upregulation of class III peroxidases promotes cross-linking of cell walls in the tissues of *Rht-B13b* carriers, constraining cell elongation and ultimately reducing height.

**Figure 7.**
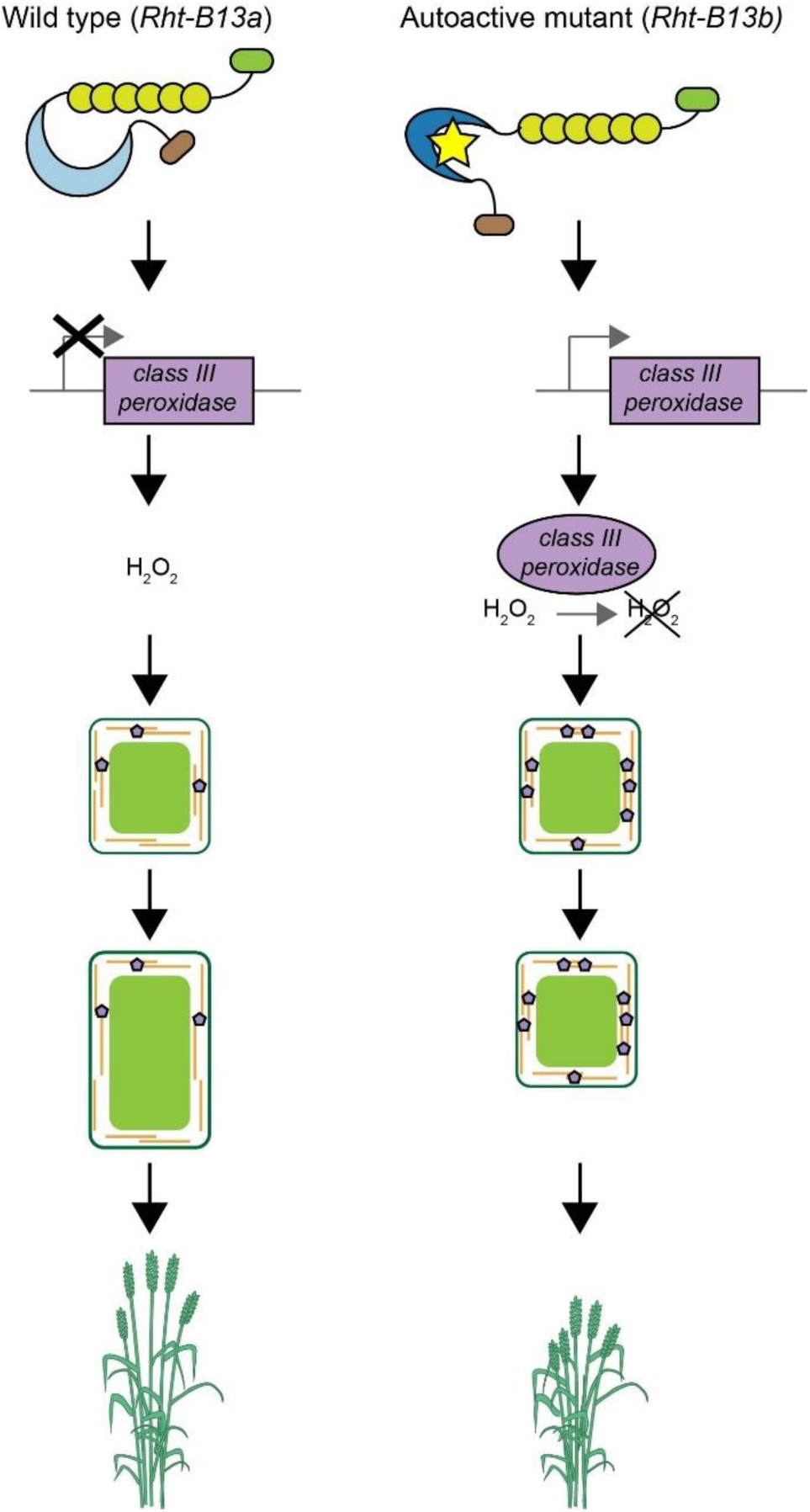
Model of pathway through which *Rht-B13b* causes semidwarfism. In a wild type plant (*Rht-B13a*, left) the NB-LRR protein is inactive resulting in normal cell wall cross-linking, cell expansion and growth. In the autoactive mutant (*Rht-B13b*, right), pathogenesis-related genes including class III peroxidases are upregulated in expanding tissues. Class III peroxidases use H_2_O_2_ to increase cell wall cross-linking which results in reduced cell expansion and growth.

### Applications in agriculture

*Rht13* is effective in multiple genetic backgrounds and provides a height reduction similar to conventional dwarfing genes *Rht-B1b* and *Rht-D1b. Rht13* dwarfism is not associated with reduced seedling growth or coleoptile length and most of the height reducing effect occurs later in development (after Zadoks stage 50) which is mainly associated with reduction in peduncle growth. Therefore, the gene is well suited to water-limiting environments that require deeper planting to access available moisture and rapid leaf area development to lower evaporative losses from the soil surface. We found that *Rht-B13b* increased bending strength which may further decrease lodging and reduce yield losses compared to conventional *Rht* genes. Deployment of *Rht-B13b* will be facilitated by the use of a perfect KASP marker for selection of the allele in breeding programmes. It is possible that *Rht-B13b* mutation is already circulating in some breeding materials, for example in the WM-800 eight-way MAGIC population of European winter wheat cultivars, a significant QTL was identified on chromosome 7B, for which the peak SNP marker maps only 10 Mb away from the location of *Rht13* (Sannemann *et al*., 2018). However, no height QTL were identified on chromosome 7B in other MAGIC populations including a diverse UK 16 founder MAGIC population (Scott *et al*., 2021) and an Australian 4 way magic population (Huang *et al*., 2012).

In conclusion, the identification of a *NB-LRR* gene underlying an alternative dwarfing gene in wheat has provided insight into an alternative pathway, where GA biosynthesis or signalling is not directly affected. This discovery will open up new opportunities to alter height, potentially through engineering of autoactive *NB-LRR* genes and cell wall enzymes. More knowledge will be needed to establish whether the activation of defense responses by *Rht-B13b* could influence disease resistance.

## Author contributions

Conceptualization: PB, WS

Data curation: PB, BF

Formal analysis: PB, WB, BF, CJP, WS, RW, SW, SJW, TX

Funding acquisition: PB, WS

Investigation: PB, DB, AD, JD, BF, JH, CM, IM, RM, RW, TX, XX

Methodology: PB, WS

Project administration: PB, WS

Resources: PB, WDB, OAO, WS

Supervision: PB, WS

Visualization: PB, WB, SW

Writing-original draft: PB

Writing-review and editing: PB, WB, WDB, AD, JD, BF, JH, IM, RM, OAO, CJP, WS, RW, SW, SJW, TX

## Data availability

The data that supports the findings of this study are available in the supplementary material of this article and raw reads for the chromosome-seq and RNA-seq are deposited as PRJEB51492 in the European Nucleotide Archive.

## Acknowledgements

We thank Zbigniew Stachurski for assistance with stem physical property measurements and Bujie Zhan for assistance with BAC libraries. We thank Jan Vrána, Zdeňka Dubská, Romana Šperková and Jitka Weiserová for assistance with chromosome sorting and DNA amplification. This research was supported by the NBI Research Computing group through HPC resources.

## References

Allan RE. 1980. Influence of semidwarfism and genetic background on stand establishment of wheat. Crop Science 20(5): 634–638.

Artimo P, Jonnalagedda M, Arnold K, Baratin D, Csardi G, de Castro E, Duvaud S, Flegel V, Fortier A, Gasteiger E, et al. 2012. ExPASy: SIB bioinformatics resource portal. Nucleic Acids Research 40(W1): W597–W603.

Bolger AM, Lohse M, Usadel B. 2014. Trimmomatic: a flexible trimmer for Illumina sequence data. Bioinformatics 30(15): 2114–2120.

Buss W, Ford BA, Foo E, Schnippenkoetter W, Borrill P, Brooks B, Ashton AR, Chandler PM, Spielmeyer W. 2020. Overgrowth mutants determine the causal role of gibberellin GA2oxidaseA13 in Rht12 dwarfism of wheat. Journal of Experimental Botany 71(22): 7171–7178.

Camacho C, Coulouris G, Avagyan V, Ma N, Papadopoulos J, Bealer K, Madden TL. 2009. BLAST+: architecture and applications. BMC Bioinformatics 10: 421.

Cavanagh CR, Chao S, Wang S, Huang BE, Stephen S, Kiani S, Forrest K, Saintenac C, Brown-Guedira GL, Akhunova A, et al. 2013. Genome-wide comparative diversity uncovers multiple targets of selection for improvement in hexaploid wheat landraces and cultivars. Proceedings of the National Academy of Sciences 110(20): 8057–8062.

Chintamanani S, Hulbert SH, Johal GS, Balint-Kurti PJ. 2010. Identification of a maize locus that modulates the hypersensitive defense response, using mutant-assisted gene identification and characterization. Genetics 184(3): 813–825.

Divashuk MG, Kroupin PY, Shirnin SY, Vukovic M, Kroupina AY, Karlov GI. 2020. Effect of gibberellin responsive reduced height allele Rht13 on agronomic traits in spring bread wheat in field experiment in non-black soil zone. Agronomy 10(7): 927.

Ellis MH, Rebetzke GJ, Azanza F, Richards RA, Spielmeyer W. 2005. Molecular mapping of gibberellin-responsive dwarfing genes in bread wheat. Theoretical and Applied Genetics 111(3): 423–430.

Ellis MH, Rebetzke GJ, Chandler P, Bonnett D, Spielmeyer W, Richards RA. 2004. The effect of different height reducing genes on the early growth of wheat. Functional Plant Biology 31(6): 583–589.

Farnham G, Baulcombe DC. 2006. Artificial evolution extends the spectrum of viruses that are targeted by a disease-resistance gene from potato. Proceedings of the National Academy of Sciences 103(49): 18828–18833.

Ford BA, Foo E, Sharwood R, Karafiátová M, Vrána J, MacMillan C, Nichols DS, Steuernagel B, Uauy C, Doležel J, et al. 2018. Rht18 semidwarfism in wheat is due to increased GA 2-oxidaseA9 expression and reduced GA content. Plant Physiology 177(1): 168–180.

Garrison E, Marth G. 2012. Haplotype-based variant detection from short-read sequencing. arXiv preprint: arXiv:1207.3907 [q-bio.GN].

Giannakopoulou A, Steele JFC, Segretin ME, Bozkurt TO, Zhou J, Robatzek S, Banfield MJ, Pais M, Kamoun S. 2015. Tomato I2 immune receptor can be engineered to confer partial resistance to the oomycete Phytophthora infestans in addition to the fungus Fusarium oxysporum. Molecular Plant-Microbe Interactions 28(12): 1316–1329.

Giorgi D, Farina A, Grosso V, Gennaro A, Ceoloni C, Lucretti S. 2013. FISHIS: fluorescence in situ hybridization in suspension and chromosome flow sorting made easy. PLOS ONE 8(2): e57994–e57994.

Haque M, Martinek P, Watanabe N, Kuboyama T. 2011. Genetic mapping of gibberellic acid-sensitive genes for semi-dwarfism in durum wheat. Cereal Research Communications 39(2): 171–178.

Hedden P. 2003. The genes of the Green Revolution. Trends in Genetics 19(1): 5–9.

Howe KL, Contreras-Moreira B, De Silva N, Maslen G, Akanni W, Allen J, Alvarez-Jarreta J, Barba M, Bolser DM, Cambell L, et al. 2020. Ensembl Genomes 2020— enabling non-vertebrate genomic research. Nucleic Acids Research 48(D1): D689–D695.

Howles P, Lawrence G, Finnegan J, McFadden H, Ayliffe M, Dodds P, Ellis J. 2005. Autoactive alleles of the flax L6 rust resistance gene induce non-race-specific rust resistance associated with the hypersensitive response. Mol Plant Microbe Interact 18(6): 570–582.

Huang BE, George AW, Forrest KL, Kilian A, Hayden MJ, Morell MK, Cavanagh CR. 2012. A multiparent advanced generation inter-cross population for genetic analysis in wheat. Plant Biotechnology Journal 10(7): 826–839.

Hyles J, Vautrin S, Pettolino F, MacMillan C, Stachurski Z, Breen J, Berges H, Wicker T, Spielmeyer W. 2017. Repeat-length variation in a wheat cellulose synthase-like gene is associated with altered tiller number and stem cell wall composition. Journal of Experimental Botany 68(7): 1519–1529.

Iwgsc, Appels R, Eversole K, Stein N, Feuillet C, Keller B, Rogers J, Pozniak CJ, Choulet F, Distelfeld A, et al. 2018. Shifting the limits in wheat research and breeding using a fully annotated reference genome. Science 361(6403): eaar7191.

Kim D, Paggi JM, Park C, Bennett C, Salzberg SL. 2019. Graph-based genome alignment and genotyping with HISAT2 and HISAT-genotype. Nature Biotechnology 37(8): 907–915.

Konzak C. 1982. Evaluation and genetic analysis of semi-dwarf mutants of wheat. In Semi-Dwarf Cereal Mutants and Their Use in Cross-Breeding: Research Coordination Meeting 1981. International Atomic Energy Agency, Vienna, Austria: 25–37.

Krasileva KV, Vasquez-Gross HA, Howell T, Bailey P, Paraiso F, Clissold L, Simmonds J, Ramirez-Gonzalez RH, Wang X, Borrill P, et al. 2017. Uncovering hidden variation in polyploid wheat. Proceedings of the National Academy of Sciences 114(6): E913–E921.

Kubaláková M, Kovárová P, Suchánková P, Cíhalíková J, Bartos J, Lucretti S, Watanabe N, Kianian SF, Dolezel J. 2005. Chromosome sorting in tetraploid wheat and its potential for genome analysis. Genetics 170(2): 823–829.

Li H, Handsaker B, Wysoker A, Fennell T, Ruan J, Homer N, Marth G, Abecasis G, Durbin R. 2009. The Sequence Alignment/Map format and SAMtools. Bioinformatics 25(16): 2078–2079.

Love MI, Huber W, Anders S. 2014. Moderated estimation of fold change and dispersion for RNA-seq data with DESeq2. Genome Biology 15(12): 550.

Mago R, Zhang P, Vautrin S, Šimková H, Bansal U, Luo M-C, Rouse M, Karaoglu H, Periyannan S, Kolmer J, et al. 2015. The wheat Sr50 gene reveals rich diversity at a cereal disease resistance locus. Nature Plants 1(12): 15186.

Marçais G, Delcher AL, Phillippy AM, Coston R, Salzberg SL, Zimin A. 2018. MUMmer4: A fast and versatile genome alignment system. PLOS Computational Biology 14(1): e1005944.

Marchler-Bauer A, Bo Y, Han L, He J, Lanczycki CJ, Lu S, Chitsaz F, Derbyshire MK, Geer RC, Gonzales NR, et al. 2017. CDD/SPARCLE: functional classification of proteins via subfamily domain architectures. Nucleic Acids Research 45(D1): D200–D203.

McIntosh RA, Dubcovsky J, Rogers WJ, Xia XC, Raupp WJ. 2020. Catalogue of Gene Symbols for Wheat. https://wheat.pw.usda.gov/GG3/WGC.

Meyers BC, Kozik A, Griego A, Kuang H, Michelmore RW. 2003. Genome-wide analysis of NBS-LRR–encoding genes in Arabidopsis. The Plant cell 15(4): 809–834.

Molnár I, Vrána J, Burešová V, Cápal P, Farkas A, Darkó É, Cseh A, Kubaláková M, Molnár-Láng M, Doležel J. 2016. Dissecting the U, M, S and C genomes of wild relatives of bread wheat (Aegilops spp.) into chromosomes and exploring their synteny with wheat. The Plant Journal 88(3): 452–467.

Moore JW, Herrera-Foessel S, Lan C, Schnippenkoetter W, Ayliffe M, Huerta-Espino J, Lillemo M, Viccars L, Milne R, Periyannan S, et al. 2015. A recently evolved hexose transporter variant confers resistance to multiple pathogens in wheat. Nature Genetics 47(12): 1494–1498.

Nakagawa T, Kurose T, Hino T, Tanaka K, Kawamukai M, Niwa Y, Toyooka K, Matsuoka K, Jinbo T, Kimura T. 2007. Development of series of gateway binary vectors, pGWBs, for realizing efficient construction of fusion genes for plant transformation. Journal of Bioscience and Bioengineering 104(1): 34–41.

Pallotta M, Warner P, Fox R, Kuchel H, Jefferies S, Langridge P 2003. Marker assisted wheat breeding in the southern region of Australia. Proceedings of the 10th international wheat genetics symposium, Paestum, Italy: Istituto Sperimentale per la Cerealicultura Roma, Italy. 789–791.

Pearce S, Huttly AK, Prosser IM, Li Y-d, Vaughan SP, Gallova B, Patil A, Coghill JA, Dubcovsky J, Hedden P, et al. 2015. Heterologous expression and transcript analysis of gibberellin biosynthetic genes of grasses reveals novel functionality in the GA3ox family. BMC Plant Biology 15(1): 130.

Peng J, Richards DE, Hartley NM, Murphy GP, Devos KM, Flintham JE, Beales J, Fish LJ, Worland AJ, Pelica F, et al. 1999. ‘Green revolution’ genes encode mutant gibberellin response modulators. Nature 400(6741): 256–261.

Pertea M, Pertea GM, Antonescu CM, Chang T-C, Mendell JT, Salzberg SL. 2015. StringTie enables improved reconstruction of a transcriptome from RNA-seq reads. Nature Biotechnology 33(3): 290–295.

Pfaffl MW. 2001. A new mathematical model for relative quantification in real-time RT–PCR. Nucleic Acids Research 29(9): e45–e45.

Pradhan Mitra P, Loqué D. 2014. Histochemical staining of Arabidopsis thaliana secondary cell wall elements. Journal of visualized experiments : JoVE(87): 51381.

Ramirez-Gonzalez RH, Segovia V, Bird N, Fenwick P, Holdgate S, Berry S, Jack P, Caccamo M, Uauy C. 2015a. RNA-Seq bulked segregant analysis enables the identification of high-resolution genetic markers for breeding in hexaploid wheat. Plant Biotechnology Journal 13(5): 613–624.

Ramirez-Gonzalez RH, Uauy C, Caccamo M. 2015b. PolyMarker: A fast polyploid primer design pipeline. Bioinformatics 31(12): 2038–2039.

Rebetzke GJ, Ellis MH, Bonnett DG, Condon AG, Falk D, Richards RA. 2011. The Rht13 dwarfing gene reduces peduncle length and plant height to increase grain number and yield of wheat. Field Crops Research 124(3): 323–331.

Rebetzke GJ, Ellis MH, Bonnett DG, Mickelson B, Condon AG, Richards RA. 2012. Height reduction and agronomic performance for selected gibberellin-responsive dwarfing genes in bread wheat (Triticum aestivum L.). Field Crops Research 126: 87–96.

Richards RA, Rebetzke GJ, Watt M, Condon AG, Spielmeyer W, Dolferus R. 2010. Breeding for improved water productivity in temperate cereals: phenotyping, quantitative trait loci, markers and the selection environment. Functional Plant Biology 37(2): 85–97.

Richardson T, Thistleton J, Higgins TJ, Howitt C, Ayliffe M. 2014. Efficient Agrobacterium transformation of elite wheat germplasm without selection. Plant Cell, Tissue and Organ Culture 119(3): 647–659.

Roberts M, Tang S, Stallmann A, Dangl JL, Bonardi V. 2013. Genetic requirements for signaling from an autoactive plant NB-LRR intracellular Innate ommune receptor. PLOS Genetics 9(4): e1003465.

Ruijter JM, Ramakers C, Hoogaars WMH, Karlen Y, Bakker O, van den Hoff MJB, Moorman AFM. 2009. Amplification efficiency: linking baseline and bias in the analysis of quantitative PCR data. Nucleic Acids Research 37(6): e45–e45.

Sannemann W, Lisker A, Maurer A, Léon J, Kazman E, Cöster H, Holzapfel J, Kempf H, Korzun V, Ebmeyer E, et al. 2018. Adaptive selection of founder segments and epistatic control of plant height in the MAGIC winter wheat population WM-800. BMC Genomics 19(1): 559.

Savelli B, Li Q, Webber M, Jemmat AM, Robitaille A, Zamocky M, Mathé C, Dunand C. 2019. RedoxiBase: A database for ROS homeostasis regulated proteins. Redox Biology 26: 101247.

Schmidt R, Kunkowska AB, Schippers JHM. 2016. Role of reactive oxygen species during cell expansion in leaves. Plant Physiology 172(4): 2098–2106.

Schopfer P. 1996. Hydrogen peroxide-mediated cell-wall stiffening in vitro in maize coleoptiles. Planta 199(1): 43–49.

Scott MF, Fradgley N, Bentley AR, Brabbs T, Corke F, Gardner KA, Horsnell R, Howell P, Ladejobi O, Mackay IJ, et al. 2021. Limited haplotype diversity underlies polygenic trait architecture across 70 years of wheat breeding. Genome Biology 22(137): 1–30.

Šimková H, Svensson JT, Condamine P, Hřibová E, Suchánková P, Bhat PR, Bartoš J, Šafář J, Close TJ, Doležel J. 2008. Coupling amplified DNA from flow-sorted chromosomes to high-density SNP mapping in barley. BMC Genomics 9(1): 294.

Smirnoff N, Arnaud D. 2019. Hydrogen peroxide metabolism and functions in plants. New Phytologist 221(3): 1197–1214.

Srinivasachary, Gosman N, Steed A, Hollins TW, Bayles R, Jennings P, Nicholson P. 2009. Semi-dwarfing Rht-B1 and Rht-D1 loci of wheat differ significantly in their influence on resistance to Fusarium head blight. Theor Appl Genet 118(4): 695–702.

Sun L, Yang W, Li Y, Shan Q, Ye X, Wang D, Yu K, Lu W, Xin P, Pei Z, et al. 2019. A wheat dominant dwarfing line with Rht12, which reduces stem cell length and affects gibberellic acid synthesis, is a 5AL terminal deletion line. The Plant Journal 97(5): 887–900.

Supek F, Bošnjak M, Škunca N, Šmuc T. 2011. REVIGO summarizes and visualizes long lists of gene ontology terms. PLOS ONE 6(7): e21800.

Talbot MJ, White RG. 2013. Cell surface and cell outline imaging in plant tissues using the backscattered electron detector in a variable pressure scanning electron microscope. Plant Methods 9(1): 40.

Tameling WIL, Vossen JH, Albrecht M, Lengauer T, Berden JA, Haring MA, Cornelissen BJC, Takken FLW. 2006. Mutations in the NB-ARC domain of I-2 that impair ATP hydrolysis cause autoactivation. Plant Physiology 140(4): 1233–1245.

Tang T. 2016. Physiological and genetic studies of an alternative semi-dwarfing gene Rht18 in wheat. University of Tasmania.

Thomas SG. 2017. Novel Rht-1 dwarfing genes: tools for wheat breeding and dissecting the function of DELLA proteins. Journal of Experimental Botany 68(3): 354–358.

Tian X, Xia X, Xu D, Liu Y, Xie L, Hassan MA, Song J, Li F, Wang D, Zhang Y, et al. 2022. Rht24b, an ancient variation of TaGA2ox-A9, reduces plant height without yield penalty in wheat. New Phytologist 233(2): 738–750.

Training W. nd. Wheat DNA extraction in 96-well plates. http://www.wheat-training.com/wp-content/uploads/Wheat_growth/pdfs/DNA_extraction_protocol.pdf.

Uauy C, Distelfeld A, Fahima T, Blechl A, Dubcovsky J. 2006. A NAC gene regulating senescence improves grain protein, zinc, and iron content in wheat. Science 314(5803): 1298–1301.

Vrána J, Kubaláková M, Simková H, Cíhalíková J, Lysák MA, Dolezel J. 2000. Flow sorting of mitotic chromosomes in common wheat (Triticum aestivum L.). Genetics 156(4): 2033–2041.

Walkowiak S, Gao L, Monat C, Haberer G, Kassa MT, Brinton J, Ramirez-Gonzalez RH, Kolodziej MC, Delorean E, Thambugala D, et al. 2020. Multiple wheat genomes reveal global variation in modern breeding. Nature 588(7837): 277–283.

Wang M, Li Z, Matthews PR, Upadhyaya NM, Waterhouse PM 1998. Improved vectors for Agrobacterium tumefaciens-mediated transformation of monocot plants: International Society for Horticultural Science (ISHS), Leuven, Belgium. 401–408.

Wang S, Wong D, Forrest K, Allen A, Chao S, Huang BE, Maccaferri M, Salvi S, Milner SG, Cattivelli L, et al. 2014. Characterization of polyploid wheat genomic diversity using a high-density 90 000 single nucleotide polymorphism array. Plant Biotechnology Journal 12(6): 787–796.

Wang Y, Chen L, Du Y, Yang Z, Condon AG, Hu Y-G. 2014. Genetic effect of dwarfing gene Rht13 compared with Rht-D1b on plant height and some agronomic traits in common wheat (Triticum aestivum L.). Field Crops Research 162: 39–47.

Wang Y, Du Y, Yang Z, Chen L, Condon AG, Hu Y-G. 2015. Comparing the effects of GA-responsive dwarfing genes Rht13 and Rht8 on plant height and some agronomic traits in common wheat. Field Crops Research 179: 35–43.

Yan J, Su P, Li W, Xiao G, Zhao Y, Ma X, Wang H, Nevo E, Kong L. 2019. Genome-wide and evolutionary analysis of the class III peroxidase gene family in wheat and Aegilops tauschii reveals that some members are involved in stress responses. BMC Genomics 20(1): 666.

Yang S, Hua J. 2004. A haplotype-specific Resistance gene regulated by BONZAI1 mediates temperature-dependent growth control in Arabidopsis. The Plant cell 16(4): 1060–1071.

Young MD, Wakefield MJ, Smyth GK, Oshlack A. 2010. Gene ontology analysis for RNA-seq: accounting for selection bias. Genome Biology 11(2): R14.

Zhang W, Chen S, Abate Z, Nirmala J, Rouse MN, Dubcovsky J. 2017. Identification and characterization of Sr13, a tetraploid wheat gene that confers resistance to the Ug99 stem rust race group. Proceedings of the National Academy of Sciences 114(45): E9483–E9492.

