## Supplementary File 1 for "An autoactive *NB-LRR* gene causes *Rht13* dwarfism in wheat"

**Supplementary file 1.** Sequence of Rht13 in CDC Stanley including genomic, CDS and protein sequence. Exons are highlighted in blue/green, UTRs in grey (predicted from RNA-seq data). The SNP mutation is highlighted in yellow. Boxes represent positions of the KASP primers and red text highlights the start and stop codon. In the protein sequence the NB-ARC domain is in blue text (e-value 9.87e-51) and the leucine rich repeat (LRR) is in green text (e-value 3.57e-06).

```
>chr7B_part2: 340124718-340118729 MSTRG.55039 (reverse complement)

CGGCGCCCGCCGGAGTGTGTCCAACGGCGGAATGTAGGTCTCCTTGAGAGTCCGTAATACTAGAATCACTTAATTGCATTATCGGTTTCGGTTC
TTCCCTTTAATACAGTACCTAAACAATATAGTGTTCTGATAGACTTTCTCGTTCGATAAATTTGCAGGTAATTCCGTGGATTCAGACTTTTTAA
TGCTCACAGTTGTCGTCCTTAGGAGACTGTTATAGCCTGCTACAGTACTGTTCTTGGAGAAGAAGCACAGTAGTAGACAGGGAACAGAGAAAAG
GGATGGGAAATTCAGACTGTATTTCATTAGCTGACCAACACACACACACACTCACACACTCACACACAAAGCCAAGCCCCGTTCCCTGATCCTA
TGTCTGCTCGTGACTGACATGAGTCACCGGCCATGACAGACAGAGACCTACTGTATTATTACTTCGCTGGGACATCAGTACATCACTTTCTCAG
TTCCTAGGACACGCACCTGTTGGACGAAACCGAGCCCACTGACGAACACAGCCAAGCAAAATTAAGGCAATGAAAAGACCTAAAGCTAACTCAT
AGGACTGGCGGGACAGCACATCTCCTCATTACTCGCCTCCACCCCGACAAAAGTCGCTACGAGGAGGCCATGGCGCTGTCCAGTCGTCCATCCT
CCTCGTTCGTCTCTGCTTCTACAGCAACCGTTGGGATCAACCAAGGACTGAGTGGGGCTCGGGTAAGACGACAGTCCACAGGCGGGGGAGGCCC
ATCCTACATTGCTCGCATCTGCTTCATCGCACAGCAGAATTTCAGTACATTCATTCTAGTGCAAGCACAGCACCAGCTTTAATTTACTGGGCCA
ATTTTCAGTACAGCTATCCATTCTAGAGGAGCCAGTTGTTAGTTGTGCCAAATAATTCAATCGCTTTTACTGCCTCGTCGTCCTCCTTTCACAT
GGGGTTTCTCCCTTTTGGAGATTGACATGACAAGGGAGACAGTGTGTCGGTTTATACTTATTGAGATTCATCATTTACTTGCGCAGCTGCATCC
TCTCTCCATTTTATCAAGCAGGGAAGAGAAAAGAAGAAGAAGAAAGCACTGCCTCCTCCTTCTCATCGTCTAAGCACCGGGATCATCCTCCTCC
GGCGCACTTCCGTCCACAGCAAGTCCCGGAATCATTCACATGTAGGTCTCCTCGAAACTCAATGCTAGCATTTTTGTTTTTGGTTCCATTTCTC
AGAAGCAAGCAAGCAATCACATTATTTGCTCACACCATTATTGTCTTCCTCACCTTGTTGCTTCTCTGTGTTCTTTGCATGTGTTTTCCTTAGT
GTTCCGATGGCGGAGCTGGTGGCCACCGTGGTGGTTGGACCACTGCTCTCCATTCTCAACGATAAGGTATCCAGCAGCCTCCTTGACCAGTACA
AGGTGATGAAAGGCATGGAGGAGCAACATGAGATCCTGATGCGTAAGCTTCCTGCCATTCTGGACATCATCGACGACGCTGAGCAGGCGGCATC
CCTGAGAAGAGGTGCAGCGGCCTGGCTTGAGGCCATCAAGAAGGTGGCTTACCAGGCCAATGAAGTCTTTGATGAGTTCAAGTACGAGGCGCTT
CGCCGCAAGGCCAAAAAGGAGGGACACTACAAGGACCTTGGCTTTGATGTGGTAAAACTCTTTCCCACCCACAACCGCTTCGTGTTCCGTAACA
GGATGGGAAGAAAGCTCCGCAAGATTGTGCAGGCCATCGAGGTCCTTGTGACCGAAATGAACGCCTTTGGCTTTAAGTATCAGCAACAAACGCC
GGTATCCAGTCAGTTGCGGCAGACGGATCCTACGATCACTGACTCGGAGGAAATCAAGAAAATCATCAATGAATCCAGAGCCAATGATAAGGAT
GAAATTGTTAGTAGACTACGTGCGCAAGCTAACAATGCAAATCTCACGGTTATTCCCATCGTTGGAATGGGCGGTCAGGGCAAGACCACCTTAG
CTCAACTAGTTTACAATGAATGTGCAGATATGAATCATTTTGATTTGCTGCTATGGGTGTGCGTCT[C/T]TGACTGCTTTGATGTGGATTCT
CTAGCTACACGTATAGTTGAAGCAGCTCGTGAGAGGAAGGATTATGGTAAAGAGGCAGCTCGTGTGAAGAAGAATGATGGTAAAGAAGCAGCT
CGTGAGAAGAAAGATGATGGTAAAGAAGCAACTCGTGAGAAGAAGGATGATAGTAAGGAAGCAGCTCCACCGAAGAAACCACTGGATTGCCTTC
AGAATGTAGTGAGCGGGCAAAGGTACCTCCTTGTGTTGGATGATGTCTGGAGACATCAGGCTAATATCTGGGATAAGCTCAAGGCTCGTCTTCA
ACATGATGGCAGCGGTAGTGTGGTCTTGATAACAACTCGTGATAAAGGACTGGTTGAAATAATGGACACTGATGAACCTCACAATCTGTCTGCT
TTGGAAGATAAATACATAAAGGAAATCATCGAGAGAAGAGCATTCAACCATTTACACAAGGAACAGGAAAGGCTCACTGGGTTGGTGAGTATGG
TTAGTGAGTTTGTAAGGAGATGTGCTGGCTCTCCTTTAGCTGCAACAGCACTGGGTTCTGTACTGCATACCAAGACCAGTAAGCAAGAATGGAT
AGATGTATTAAGCAAAAGCAGCATTTGCACCAAGGAATCTGGAATCTTACCAATACTCAAGCTCAGTTACAGCGACTTGCCGTCGCATATGAAA
CCATGCTTTGCTTTTTGTGCTGTATTTCCTAAAGATTATGAAATTGATGTGGACAAGCTGATCCAACTATGGATTGCACATGGCTTCATCCATG
AAAAGCAAGGTCATCTTGAAACCATTGGCAAAGCGATTTTCCATGAGTTGGCCTCAAGGTCTTTCTTTCAGGATGTGGAACAAGTCCAAGCCAC
AAGTAGTCAGCAATCCATGTTGTGTTACTCTAGAACAACATGTAAAATCCATGATCTTATGCATGATGTTGCACTTTCGGTAATGGAAAAGGAA
TGCGCCTTGGCAACTGAGGAACCAGGCAAGATTGAATCTGCTGTCGCAACTGAGGAACCAAGTCAGAGTGAGTGGCTTCCAAACACAGCTCGGC
ATTTATTTTTGTCATGCAAGGACCCAGAAAAAAAATTGAATAGTTCTCTGGAGAACAGCTTTCCAGCCATCCAAACACTTCTGTGTGATAGATA
TATGAGTAGTTCATTGCAGCATCTATCAAAGTACAGCTCTCTGCAAGCATTACAGCTCCATTTACTTAGAAGATCATTTCCATTGAAACCAAAG
TATCTACATCACCTGAGGTACCTAGATCTTTCTAGAAGTTGGATCAAAGCACTTCCCGAAGATATGAGCATTCTACACAACCTGCAAACGCTTA
ACCTTTCTGGATGTGAATATCTTGAAACACTTCCGAGACAAATGAAGTATATGATAGCCCTGCGTCACCTTTACACTCATGGTTGTCCAGAGCT
GAAGAGCATGCCAAGAGACCTCGGAAAACTCACATCCCTACAGACACTTACATGCTTTGTAGCAGCTAATAGCTCTAGTTGCAGTAATGTGGGA
CAGATTGGGAATCTAAAACTTGGTGGTCAACTAGAGCTACGTAATCTGGCAAATGTGACAGAAGTGGATGCAGAAGCAGCAAATCTCATGAACA
AGGAGGAGCTAAGAAAACTGACATTAACATGGACCTTGGGATGGAATTATTCTGAAGATAAAACTTGCTGGCGGGATAATGAAGAGGATGCAAG
AGTGCTCAACAATCTCAAACCTCATGATGGACTAGATGCCGTAGGGATACACTCATATGGAGCCACCACCTTCCCGACATGGATGACTATGTTG
CAAAACATTGTTGAGATCCATCTTTTTGGTTGTAGAAAACTGCAATGGTTTTTCAGCCGTGACTGTGATAATAAAAGCTTTGCATTTCGAAAAC
TAAAGGAGCTTACGTTGCACGCTCTTGTCTCTTTGGAGAGATTGTGGGAGATAGATAATGATGAGATGCACAAAGAAGAAATCTTTCCTCTGCT
TGAGAAGTTGTCCATTAGTCACTGTGAAAATTTGAAAGCATTGCCAGGACAGCCGACCTTCCCTAAGCTTCAGAATGTTCGTATTAAGAAATGT
CCACAGTTGACAAGTACAGCTAAATCACCAAAGCTCAGTGTATTAAAAATGGAAGGAACTGAGATAGAGTTGTTCTTGTGGGTAGCGAGACATA
TGACTTCATTGACCAATCTGGAACTGACTAGCATTGAACATGGAACTGATACAACCTCGATGGGGGCTGAGAATAGTCTGAGGGAAGTGGTGAG
TGTCAAGGAAAAAGGGAAAGATCAAGATTTCCCTCTAGCAGTTTTGGTGTTAAGAGACTTTAAGTCAGGTGTAAGTGTACCAGATATGTGTGCA
TGCTTTGTACACCTTCAAGAGTTGTCAATTTTGGGTTGCCATGCGCTCGTCCACTGGCCAGAAAAATTGTTTGAAGGATTGGTATTCTTGAGGA
GGCTTCATATTGCAGATTGTGATAATCTGACTGGATATGCACAAGCTTCTGCTGAGCCATCAACGTCATCAGAAACCGGTCAGCTCCTGCCACG
TCTAAAGTCTCTGTCGATAATGAGTTGTGAAAACTTGGTTGAGCTCTTCAACGTCCCTGCATCTCTCAGGAAAATTTACATTTATAATTGCAAT
AAGCTCGAGACCACATGCGGCAGGAAGCAGCAGCAGGGACAGTCAGTATCATCGACTCATCAAGGGTCATCCAGTATAGAAGAATTATCATTCT
ACACCTGTCATGGCTTAACAGGGGTCCTCTATATTCCCCCATCCCTGAAGAAACTAGACATTTTCAACTGTCGTGGGTTGGCATCTCTGGAATC
CTGCTCTCCCGAGCTCCAATCCTTGGAGTACCTCCAGCTTGGGCACTGCAATTCCCTGTCATCCCTACCGGATGTGCCGCAAGCATACTCATCT
CTCCAATATCTTACCATTAACGACTGCCCTGGCATGAAGATGCTCCCTGCAAGCCTGCAACAACGCCTGGGCAGCATCCAACATGTGTACATAG
ATGCCCATTATTATGGAAGTAAGCACACATCAATTACCTTCTCTATTTCTACTTACTAGCGAGTGTTAAATGTGCTAGTCAGATAATGATAATA
GCATCACTTGTAGGACGACAGAAAACTGGAGCAAGTCAGATATTCCTTTATCTAAATCCATCATTTCTTTCTTTAATGATGTTCTTCGAACTGT
AAGTTTTGCATGACTAATCTGTGTAGATAAGCCTATGCTGCTGAAGCCGAAGACATGGAAGTATGTCTGCAAAGGTTGAACTACTGTTCCAGAG
```

GTGTTGTCTGCAAAGGTTGGTGGCACTGACTGTACCCCTCGGATATATTTTGAGGAAAAAAATACGTGCCACATCATTATGTATTTGGCTTTCT
TCTGGTGTTGACCGTGTATGGTTTCTTTCTTCAATTCTTCCAGCACGTGCATACGGGTGGAAAAGAGAGATCCCTTGGAGTAGAACTAGTGTCT
CGCAAGACCTGCCAGGATCAGCACCTGTTGCCTCCATGTTCGTTTTGTGATGTTGCCTGGACCAGACGCCGCCATGAATGTTGTTGCCATGTTG
AGTTGCAATGTCAGTAATCAGAAAGTGCTTCAAACACTATTGTTTTATCCCCGGGAAGAGATCTTGTTTGAACAATTGTATTTTCCACTTGTTT
ACCTATGGGCTGTGATATGATTGTACTATTTATTTTTTGAACTGGCAAGGCGACTCTCTGCCCCTGGCACTGTCATTATTATTTTTTTTGTCTA
AATTTTGCCCAGAAGAGGATACTTGTTGACTCAGTTTTTTGTATGTATGTCCGTCAGTCTCTGTATGACTTGTGTGTAACTTGGAAGCTCAACG
TTTGCTATATATTTTGGTTTGTAAACACATGCATTGTTATTTAGTGATCATCATATACACATCTGAAGTTGTT

>MSTRG.55039 CDS
ATGGCGGAGCTGGTGGCCACCGTGGTGGTTGGACCACTGCTCTCCATTCTCAACGATAAGGTATCCAGCAGCCTCCTTGACCAGTACAAGGTGA
TGAAAGGCATGGAGGAGCAACATGAGATCCTGATGCGTAAGCTTCCTGCCATTCTGGACATCATCGACGACGCTGAGCAGGCGGCATCCCTGAG
AAGAGGTGCAGCGGCCTGGCTTGAGGCCATCAAGAAGGTGGCTTACCAGGCCAATGAAGTCTTTGATGAGTTCAAGTACGAGGCGCTTCGCCGC
AAGGCCAAAAAGGAGGGACACTACAAGGACCTTGGCTTTGATGTGGTAAAACTCTTTCCCACCCACAACCGCTTCGTGTTCCGTAACAGGATGG
GAAGAAAGCTCCGCAAGATTGTGCAGGCCATCGAGGTCCTTGTGACCGAAATGAACGCCTTTGGCTTTAAGTATCAGCAACAAACGCCGGTATC
CAGTCAGTTGCGGCAGACGGATCCTACGATCACTGACTCGGAGGAAATCAAGAAAATCATCAATGAATCCAGAGCCAATGATAAGGATGAAATT
GTTAGTAGACTACGTGCGCAAGCTAACAATGCAAATCTCACGGTTATTCCCATCGTTGGAATGGGCGGTCAGGGCAAGACCACCTTAGCTCAAC
TAGTTTACAATGAATGTGCAGATATGAATCATTTTGATTTGCTGCTATGGGTGTGCGTCT[C/T]TGACTGCTTTGATGTGGATTCTCTAGCT
ACACGTATAGTTGAAGCAGCTCGTGAGAGGAAGGATTATGGTAAAGAGGCAGCTCGTGTGAAGAAGAATGATGGTAAAGAAGCAGCTCGTGAG
AAGAAAGATGATGGTAAAGAAGCAACTCGTGAGAAGAAGGATGATAGTAAGGAAGCAGCTCCACCGAAGAAACCACTGGATTGCCTTCAGAATG
TAGTGAGCGGGCAAAGGTACCTCCTTGTGTTGGATGATGTCTGGAGACATCAGGCTAATATCTGGGATAAGCTCAAGGCTCGTCTTCAACATGA
TGGCAGCGGTAGTGTGGTCTTGATAACAACTCGTGATAAAGGACTGGTTGAAATAATGGACACTGATGAACCTCACAATCTGTCTGCTTTGGAA
GATAAATACATAAAGGAAATCATCGAGAGAAGAGCATTCAACCATTTACACAAGGAACAGGAAAGGCTCACTGGGTTGGTGAGTATGGTTAGTG
AGTTTGTAAGGAGATGTGCTGGCTCTCCTTTAGCTGCAACAGCACTGGGTTCTGTACTGCATACCAAGACCAGTAAGCAAGAATGGATAGATGT
ATTAAGCAAAAGCAGCATTTGCACCAAGGAATCTGGAATCTTACCAATACTCAAGCTCAGTTACAGCGACTTGCCGTCGCATATGAAACCATGC
TTTGCTTTTTGTGCTGTATTTCCTAAAGATTATGAAATTGATGTGGACAAGCTGATCCAACTATGGATTGCACATGGCTTCATCCATGAAAAGC
AAGGTCATCTTGAAACCATTGGCAAAGCGATTTTCCATGAGTTGGCCTCAAGGTCTTTCTTTCAGGATGTGGAACAAGTCCAAGCCACAAGTAG
TCAGCAATCCATGTTGTGTTACTCTAGAACAACATGTAAAATCCATGATCTTATGCATGATGTTGCACTTTCGGTAATGGAAAAGGAATGCGCC
TTGGCAACTGAGGAACCAGGCAAGATTGAATCTGCTGTCGCAACTGAGGAACCAAGTCAGAGTGAGTGGCTTCCAAACACAGCTCGGCATTTAT
TTTTGTCATGCAAGGACCCAGAAAAAAAATTGAATAGTTCTCTGGAGAACAGCTTTCCAGCCATCCAAACACTTCTGTGTGATAGATATATGAG
TAGTTCATTGCAGCATCTATCAAAGTACAGCTCTCTGCAAGCATTACAGCTCCATTTACTTAGAAGATCATTTCCATTGAAACCAAAGTATCTA
CATCACCTGAGGTACCTAGATCTTTCTAGAAGTTGGATCAAAGCACTTCCCGAAGATATGAGCATTCTACACAACCTGCAAACGCTTAACCTTT
CTGGATGTGAATATCTTGAAACACTTCCGAGACAAATGAAGTATATGATAGCCCTGCGTCACCTTTACACTCATGGTTGTCCAGAGCTGAAGAG
CATGCCAAGAGACCTCGGAAAACTCACATCCCTACAGACACTTACATGCTTTGTAGCAGCTAATAGCTCTAGTTGCAGTAATGTGGACAGATT
GGGAATCTAAAACTTGGTGGTCAACTAGAGCTACGTAATCTGGCAAATGTGACAGAAGTGGATGCAGAAGCAGCAAATCTCATGAACAAGGAGG
AGCTAAGAAAACTGACATTAACATGGACCTTGGGATGGAATTATTCTGAAGATAAAACTTGCTGGCGGGATAATGAAGAGGATGCAAGAGTGCT
CAACAATCTCAAACCTCATGATGGACTAGATGCCGTAGGGATACACTCATATGGAGCCACCACCTTCCCGACATGGATGACTATGTTGCAAAAC
ATTGTTGAGATCCATCTTTTTGGTTGTAGAAAACTGCAATGGTTTTTCAGCCGTGACTGTGATAATAAAAGCTTTGCATTTCGAAAACTAAAGG
AGCTTACGTTGCACGCTCTTGTCTCTTTGGAGAGATTGTGGGAGATAGATAATGATGAGATGCACAAAGAAGAAATCTTTCCTCTGCTTGAGAA
GTTGTCCATTAGTCACTGTGAAAATTTGAAAGCATTGCCAGGACAGCCGACCTTCCCTAAGCTTCAGAATGTTCGTATTAAGAAATGTCCACAG
TTGACAAGTACAGCTAAATCACCAAAGCTCAGTGTATTAAAAATGGAAGGAACTGAGATAGAGTTGTTCTTGTGGGTAGCGAGACATATGACTT
CATTGACCAATCTGGAACTGACTAGCATTGAACATGGAACTGATACAACCTCGATGGGGGCTGAGAATAGTCTGAGGGAAGTGGTGAGTGTCAA
GGAAAAAGGGAAAGATCAAGATTTCCCTCTAGCAGTTTTGGTGTTAAGAGACTTTAAGTCAGGTGTAAGTGTACCAGATATGTGTGCATGCTTT
GTACACCTTCAAGAGTTGTCAATTTTGGGTTGCCATGCGCTCGTCCACTGGCCAGAAAAATTGTTTGAAGGATTGGTATTCTTGAGGAGGCTTC
ATATTGCAGATTGTGATAATCTGACTGGATATGCACAAGCTTCTGCTGAGCCATCAACGTCATCAGAAACCGGTCAGCTCCTGCCACGTCTAAA
GTCTCTGTCGATAATGAGTTGTGAAAACTTGGTTGAGCTCTTCAACGTCCCTGCATCTCTCAGGAAAATTTACATTTATAATTGCAATAAGCTC
GAGACCACATGCGGCAGGAAGCAGCAGCAGGGACAGTCAGTATCATCGACTCATCAAGGGTCATCCAGTATAGAAGAATTATCATTCTACACCT
GTCATGGCTTAACAGGGGTCCTCTATATTCCCCCATCCCTGAAGAAACTAGACATTTTCAACTGTCGTGGGTTGGCATCTCTGGAATCCTGCTC
TCCCGAGCTCCAATCCTTGGAGTACCTCCAGCTTGGGCACTGCAATTCCCTGTCATCCCTACCGGATGTGCCGCAAGCATACTCATCTCTCCAA
TATCTTACCATTAACGACTGCCCTGGCATGAAGATGCTCCCTGCAAGCCTGCAACAACGCCTGGGCAGCATCCAACATGTGTACATAGATGCCC
ATTATTATGGAAATAAGCCTATGCTGCTGAAGCCGAAGACATGGAAGTATGTCTGCAAAGGTTGA

>WT_Protein

MAELVATVVVGPLLSILNDKVSSSLLDQYKVMKGMEEQHEILMRKLPAILDIIDDAEQAASLRRGAAAWLEAIKKVAYQANEVFDEFKYEALRR
KAKKEGHYKDLGFDVVKLFPTHNRFVFRNRMGRKLRKIVQAIEVLVTEMNAFGFKYQQQTPVSSQLRQTDPTITDSEEIKKIINESRANDKDEI
VSRLRAQANNANLTVIPIVGMGGQGKTTLAQLVYNECADMNHFDLLLWVCV[S/F]DCFDVDSLATRIVEAARERKDYGKEAARVKKNDGKEAA
REKKDDGKEATREKKDDSKEAAPPKKPLDCLQNVVSGQRYLLVLDDVWRHQANIWDKLKARLQHDGSGSVVLITTRDKGLVEIMDTDEPHNLSA
LEDKYIKEIIERRAFNHLHKEQERLTGLVSMVSEFVRRCAGSPLAATALGSVLHTKTSKQEWIDVLSKSSICTKESGILPILKLSYSDLPSHMK
PCFAFCAVFPKDYEIDVDKLIQLWIAHGFIHEKQGHLETIGKAIFHELASRSFFQDVEQVQATSSQQSMLCYSRTTCKIHDLMHDVALSVMEKE
CALATEEPGKIESAVATEEPSQSEWLPNTARHLFLSCKDPEKKLNSSLENSFPAIQTLLCDRYMSSSLQHLSKYSSLQALQLHLLRRSFPLKPK
YLHHLRYLDLSRSWIKALPEDMSILHNLQTLNLSGCEYLETLPRQMKYMIALRHLYTHGCPELKSMPRDLGKLTSLQTLTCFVAANSSSCSNVG
QIGNLKLGGQLELRNLANVTEVDAEAANLMNKEELRKLTLTWTLGWNYSEDKTCWRDNEEDARVLNNLKPHDGLDAVGIHSYGATTFPTWMTML
QNIVEIHLFGCRKLQWFFSRDCDNKSFAFRKLKELTLHALVSLERLWEIDNDEMHKEEIFPLLEKLSISHCENLKALPGQPTFPKLQNVRIKKC
PQLTSTAKSPKLSVLKMEGTEIELFLWVARHMTSLTNLELTSIEHGTDTTSMGAENSLREVVSVKEKGKDQDFPLAVLVLRDFKSGVSVPDMCA
CFVHLQELSILGCHALVHWPEKLFEGLVFLRRLHIADCDNLTGYAQASAEPSTSSETGQLLPRLKSLSIMSCENLVELFNVPASLRKIYIYNCN
KLETTCGRKQQQGQSVSSTHQGSSSIEELSFYTCHGLTGVLYIPPSLKKLDIFNCRGLASLESCSPELQSLEYLQLGHCNSLSSLPDVPQAYSS
LQYLTINDCPGMKMLPASLQQRLGSIQHVYIDAHYYGNKPMLLKPKTWKYVCKG-
