## Supplementary Figures and Tables for "An autoactive *NB-LRR* gene causes *Rht13* dwarfism in wheat"

### Supplementary Tables and Figures

### Supplementary Tables

**Supplementary Table 1.** Primers used for genetic mapping in Magnif x Magnif M population. BAC sequence-derived markers *7J15.144I10_2_2* and *127M17.134P08_3* are highlighted in red, these markers flanked the *Rht13* locus on the proximal and the distal side, respectively.

| Marker | Marker type | Primer name | Primer sequence | Co-segregating | Remarks |
|---|---|---|---|---|---|
| **gwm577** | SSR | gwm577 F | ATGGCATAATTTGGTGAAATTG | No | |
| | | gwm577 R | TGTTTCAAGCCCAACTTCTATT | | |
| **2825** | KASP | 2825 FAM | GAAGGTGACCAAGTTCATGCTGACATAACAGAGTGTCGACCTT | No | Dominant marker |
| | | 2825 VIC | GAAGGTCGGAGTCAACGGATTGACATAACAGAGTGTCGACCTG | | |
| | | 2825 com | TAGCCCAGTTCTGGCTTTGA | | |
| **3513** | KASP | 3513 FAM | GAAGGTGACCAAGTTCATGCTAGACATAACAATCCAAGCTGGT | No | Dominant marker |
| | | 3513 VIC | GAAGGTCGGAGTCAACGGATTAGACATAACAATCCAAGCTGGC | | |
| | | 3513 com | AATCTCAATGCCGGGTGTT | | |
| **7261** | KASP | 7261 FAM | GAAGGTGACCAAGTTCATGCTTCAACTGCAACGACTGAACGGA | No | Dominant marker |
| | | 7261 VIC | GAAGGTCGGAGTCAACGGATTTCAACTGCAACGACTGAACGGG | | |
| | | 7261 com | CCTCCCCTCATCAGGAAAT | | |
| **30130** | PCR | 30130 F | TGGCACAGAATTGATCTGCTT | No | Dominant marker |
| | | 30130 R | TATCCTGCTCCTGACACTCCA | | |

| | | | | | |
|---|---|---|---|---|---|
| **30150** | PCR | 30150 F2<br>30150 R2 | TGTGATCTGCACTTGCTCTTG<br>CAAGTTTCTGATCTCATCAGCC | No | Dominant marker |
| **5252** | PCR | 5252 F3<br>5252 R5 | CCTCCGTGTTACAGGTGCTT<br>GATTGGGATCAAGACATCT | No | Dominant marker |
| **6818** | KASP | 6818 FAM<br>6818 VIC<br>6818 com | GAAGGTGACCAAGTTCATGCTCAACTATTTCTCCGTGGTCA<br>GAAGGTCGGAGTCAACGGATTCAACTATTTCTCCGTGGTCT<br>ACAGTAATTTACGAACCGCA | No | |
| **159N02**<br>(159N02_17_1) | KASP | 159N02_17_1 FAM2<br>159N02_17_1 VIC2<br>159N02_17_1 com2 | GAAGGTGACCAAGTTCATGCTCGGTGACAACGGAGACACTT<br>GAAGGTCGGAGTCAACGGATTCGGTGACAACGGAGACACTG<br>GTAAGTTGGACGGTTTGGCC | No | |
| **30170 (2)** | KASP | 30170 FAM2<br>30170 VIC2<br>30170 com2 | GAAGGTGACCAAGTTCATGCTGCCAAGGGGTAGTAGTCTTATGCAG<br>GAAGGTCGGAGTCAACGGATTGCCAAGGGGTAGTAGTCTTATGCAA<br>TCGGTGCCTCTATCAAAGTG | No | |
| **30160** | PCR | 30160 F15<br>30160 R15 | CCGTGGCTCAGGAATTAAGC<br>ACTAACACACTGACGAGGCA | No | Dominant marker |
| **144I10_2** | PCR | 144I10_2 F1<br>144I10_2 R1 | GCAACGATCCTCTGATGCAG<br>AGCGAATTACCCTGCACCTA | No | Dominant marker |
| **30180** | PCR | 30180 F<br>30180 R | GTGGTTCCATATATGCTGCTG<br>GCTCCAACTGCTGTCTAGAATA | No | Dominant marker |
| **7J15**<br>(7J15.144I10_2_2) | KASP | 7J15 FAM1<br>7J17 VIC1<br>7J15 com1 | GAAGGTGACCAAGTTCATGCTCATCCGCTTAAGGTTCCTGAC<br>GAAGGTCGGAGTCAACGGATTCATCCGCTTAAGGTTCCTGAT<br>GCGGAGGATATCATAAGCGA | No | |

| | | | | | |
|---|---|---|---|---|---|
| **0521** | PCR | 0521 F1<br>0521 R1 | GGATTTCCGTTTTGAGCATCAG<br>ACGAGTTGTTGTCAGGACCAC | No | Dominant marker |
| **1323** | PCR | 1323 F4<br>1323 R4 | AGGACATCGCACGCCTTATA<br>AGACAACTGCTCAGACCGGT | No | Dominant marker |
| **138K15_1_2** | PCR | 138K15_1_2 F2<br>138K15_1_2 R2 | GGAAGATCGTGTTAATGAGATGTG<br>TATCGTCGCGGAGTTTAGGC | No | Dominant marker |
| **10J24.31N23_5** | PCR | 10J24.31N23_5 F1<br>10J24.31N23_5 R1 | TCTTTTGTGAGCCCAGCGAT<br>TGTTGGATGCTTGTTGTGCG | No | Dominant marker |
| **30190** | PCR | 30190 F1<br>30190 R1 | TGATTGAGCCTAACACAAACC<br>GCAGGCTCCGCACCTTCTCAC | Yes | Dominant marker |
| **30190** | KASP | 30190 FAM<br>30190 VIC<br>30190 com | GAAGGTGACCAAGTTCATGCTAGGACTGAATCAAGCATCTCATA<br>GAAGGTCGGAGTCAACGGATTAGGACTGAATCAAGCATCTCATC<br>GTCGTCGACTCCTCCATCC | Yes | |
| **2594** | PCR | 2594 F6<br>2594 R6 | TGAGATGGTTGGTGCGGATA<br>GGCCTTGGCTTCGTTTGTAA | Yes | Dominant marker |
| **2594** | KASP | 2594 FAM1<br>2594 VIC1<br>2594 com1 | GAAGGTGACCAAGTTCATGCTAGGAAGACGCAGGAGTACA<br>GAAGGTCGGAGTCAACGGATTAGGAAGACGCAGGAGTACG<br>GATGGGGCTCTTCACCTTG | Yes | |
| **150J11.171J01_10** | PCR | 150J11.171J01_10 F1<br>150J11.171J01_10 R1 | GCCCCTCTTGTTCTCATGCT<br>GAGACTGGCTGGCATCCTTT | Yes | Dominant marker |
| **56C12_1_1** | PCR | 56C12_1 F4<br>56C12_1 R4 | GACCGCTTCTCTCCATTGTG<br>CATGTTGAAGTCGGGCCTTA | Yes | Dominant marker |

| | | | | | |
|---|---|---|---|---|---|
| **103L05_1_1** | PCR | 103L05_1 F4<br>103L05_1 R4 | TGATTACTCGCCTCCATCCC<br>AGAATGGAGAGCAGTGGTCC | Yes | Dominant marker |
| **55D09_1** | PCR | 55D09_1 F1<br>55D09_1 R1 | GGATGAGATTGTCCGTTGGC<br>GATGCAGCGGTACAACTCTG | Yes | Dominant marker |
| **38E21.55D09_16** | PCR | 38E21.55D09_16 F4<br>38E21.55D09_16 R4 | CTACGGTGTGGTTTTGGAGC<br>GCATGCATAGACACAACGGG | Yes | Dominant marker |
| **19C10**<br>(19C10.38E21_3) | KASP | 19C10 FAM<br>19C10 VIC<br>19C10 com | GAAGGTGACCAAGTTCATGCTACTCCTCTGGATTGAAAGTGA<br>GAAGGTCGGAGTCAACGGATTACTCCTCTGGATTGAAAGTGG<br>AAGTGCTTTTCTCATGTCGC | Yes | |
| **127M17**<br>(127M17.134P08_3) | KASP | 127M17 FAM<br>127M17 VIC<br>127M17 com | GAAGGTGACCAAGTTCATGCTCCATTCAACTCAGCAACTCACC<br>GAAGGTCGGAGTCAACGGATTCCATTCAACTCAGCAACTCACT<br>ATACAGTCGTGGCCAGAAAT | No | |
| **26P16_7** | Sequencing | 26P16_7 F3<br>26P16_7 R3 | GATGGCGGTGGTGATGATAATGA<br>GGCGCAAAATCTAGGTATAAAGTC | No | |
| **182N11.187A12_1** | PCR | 182N11.187A12_1 F3<br>182N11.187A12_1 R3 | GGAAGAACTAGGGACGTAATGG<br>GGGATGGTCGAATTTGGGGT | No | Dominant marker |
| **182N11_2_1** | PCR | 182N11_2_1 F2<br>182N11_2_1 R2 | TGTCTCCGGTTGCTCATTC<br>AGGTGGATTGCATGGCGTTA | No | Dominant marker |
| **182N11 2**<br>(182N11_2_2) | KASP | 182N11 FAM2<br>182N11 VIC2<br>182N11 com2 | GAAGGTGACCAAGTTCATGCTCGTCTCATCATCTGGCCATAT<br>GAAGGTCGGAGTCAACGGATTCGTCTCATCATCTGGCCATAC<br>CTAAGAGCACCATGGAGAGC | No | |

| | | | | | |
|---|---|---|---|---|---|
| **30200** | PCR | 30200 F<br>30200 R | TTCATGACGGAGATTGATAGA<br>AATATGGATGGTTCTCTGCTC | No | Dominant marker |
| **wmc276** | SSR | | GACATGTGCACCAGAATAGC<br>AGAAGAACTATTCGACTCCT | No | |

**Supplementary Table 2.** Recombinants identified across the *Rht13* mapping interval. Markers in red correspond to BAC sequence-derived markers *7J15.144I10_2_2* (7J15) and *127M17.134P08_3* (127M17) which markers flanked the *Rht13* locus on the proximal and the distal side, respectively.

| Marker | origin | 4D1 | 6D4 | 3G3 | 5F11 | 4D11 | 6B1 | 3B6 | 11G3 | 7E5 | 41G2 | 9A3 | 12A9 | 22E4 | 30E2 | 16A4 | 8G1 | 5A10 | 5D8 | 5G3 | 6D7 | 7D8 | ML45-S | ML80-T |
|---|---|---|---|---|---|---|---|---|---|---|---|---|---|---|---|---|---|---|---|---|---|---|---|---|
| gwm577 | SSR | B | A | A | A | A | B | B | | B | | | | | | | | A | A | A | A | A | B | A |
| 2825 | SSR | B | A | A | A | A | B | B | | B | | | | | | | | A | A | A | A | A | B | A |
| 3513 | SSR | B | A | A | A | A | B | B | | B | | | | | | | | A | A | A | A | A | B | A |
| 7261 | SSR | | A | A | A | A | B | B | | B | | | | | | | | A | A | A | A | A | B | A |
| 30130 | BAC | B | A | A | A | A | B | B | | B | | | | | | | | A | A | A | A | A | B | A |
| 30150 | BAC | A | B | B | A | A | B | B | B | B | B | A | A | A | B | B | A | A | A | A | A | A | B | A |
| 6818 | BAC | A | B | B | B | A | B | B | | B | | | | | | | | A | A | A | A | A | B | A |
| 159N02 | BAC | A | B | B | B | B | B | B | | B | | | | | | | | A | A | A | A | A | B | A |
| 30170 (2) | BAC | A | B | B | B | B | A | B | B | B | B | A | A | A | B | B | A | A | A | A | A | A | B | A |
| 30160 | BAC | A | B | B | B | B | A | A | A | A | A | B | B | A | B | B | A | A | A | A | A | A | B | A |
| 144I10_2 | BAC | A | B | B | B | B | A | A | A | A | | B | B | A | B | B | A | A | A | A | A | A | B | A |
| 30180 | BAC | A | B | B | B | B | A | A | A | A | | B | B | A | B | B | A | A | A | A | A | A | B | A |
| 7J15 | BAC | A | B | B | B | B | A | A | A | A | A | B | B | A | B | B | A | A | A | A | A | A | B | A |
| 0521 | BAC | A | B | B | B | B | A | A | A | A | | B | B | B | B | B | A | A | A | A | A | A | B | A |
| 1323 | BAC | A | B | B | B | B | A | A | A | A | | B | B | B | B | B | A | A | A | A | A | A | B | A |
| 138K15_1_2 | BAC | A | B | B | B | B | A | A | A | A | A | B | B | B | B | B | A | A | A | A | A | A | B | A |

| | | | | | | | | | | | | | | | | | | | | | | | | |
|---|---|---|---|---|---|---|---|---|---|---|---|---|---|---|---|---|---|---|---|---|---|---|---|---|
| 10J24.31N23_5 | BAC | A | B | B | B | B | A | A | A | A | A | B | B | B | B | B | A | A | A | A | A | A | B | A |
| 30190 | BAC | A | B | B | B | B | A | A | A | A | A | B | B | B | A | B | A | A | A | A | A | A | B | A |
| 2594 | BAC | A | B | B | B | B | A | A | | A | | | | | | | | A | A | A | A | A | B | A |
| 150J11.171J01_10 | BAC | A | B | B | B | B | A | A | A | A | A | B | B | B | A | B | A | A | A | A | A | A | B | A |
| 56C12_1_1 | BAC | A | B | B | B | B | A | A | A | A | A | B | B | B | A | B | A | A | A | A | A | A | B | A |
| **Rht-13** | | **A** | **B** | **B** | **B** | **B** | **A** | **A** | **A** | **A** | **A** | **B** | **B** | **B** | **A** | **B** | **A** | **A** | **A** | **A** | **A** | **a** | **B** | **A** |
| 103L05_1_1 | BAC | A | B | B | B | B | A | A | A | A | A | B | B | B | A | B | A | A | A | A | A | A | B | A |
| 55D09_1 | BAC | A | B | B | B | B | A | A | A | A | A | B | B | B | A | B | A | A | A | A | A | A | B | A |
| 38E21.55D09_16 | BAC | A | B | B | B | B | A | A | A | A | A | B | B | B | A | B | A | A | A | A | A | A | B | A |
| 19C10 | BAC | A | B | B | B | B | A | A | A | A | A | B | B | B | A | B | A | A | A | A | A | A | B | A |
| 127M17 | BAC | A | B | B | B | B | A | A | A | A | A | B | B | B | A | A | A | A | A | A | A | A | B | A |
| 26P16_7 | BAC | A | B | B | B | B | A | A | A | A | A | B | B | B | A | A | B | A | A | A | A | A | B | A |
| 182N11_2_1 | BAC | A | B | B | B | B | A | A | A | A | A | B | B | B | A | A | B | A | A | A | A | A | B | A |
| 182N11 (1) | BAC | A | B | B | B | B | A | A | A | A | A | B | B | B | A | A | B | A | A | A | A | A | B | A |
| 182N11 (2) | BAC | A | B | B | B | B | A | A | | A | | | | | | | | B | A | A | A | A | B | A |
| 30200 | BAC | A | B | B | B | B | A | A | A | A | A | B | B | B | A | A | B | B | B | B | B | B | B | A |
| wmc276 | SSR | A | B | B | B | B | A | A | | A | | | | | | | | B | B | B | B | B | B | A |

**Supplementary Table 3**

Chromosome content of the 7B chromosomal samples flow sorted from short and tall progeny of the Magnif x Magnif M cross. The purity of the 7B chromosomes and the contamination by non-target chromosomes were determined by microscopic analysis of the flow sorted chromosomes after FISH.

| Genotype | Chr. | Purity of 7B (%) | Contamination (Chr.: %) | Phenotype | DNA amount after amplification (ug) | Sequencing |
|---|---|---|---|---|---|---|
| M136-6 | 7B | 80.0% | 2B: 12.59%<br>4A: 2.22%<br>5B:0.74%<br>1B:0.74% | short | 5.4 | Illumina 50 Gb |
| M103-4 | 7B | 76.78% | 2B: 18.75%<br>3B:0.89%<br>4A: 1.78%<br>5B: 0.89%<br>1B: 0.89% | short | 7.39 | Illumina 50 Gb |
| M212-9 | 7B | 81.55% | 2B: 14.56%<br>3B: 1.94%<br>5B: 0.97%<br>6B: 0.97% | short | 6.79 | Illumina 50 Gb |
| M286-4 | 7B | 82.25% | 2B: 14.51%<br>6B: 1.61%<br>1B: 1.61% | short | 7.91 | Illumina 50 Gb |
| M157-4 | 7B | 70.0% | 2B: 27.64%<br>3B:2.35% | tall | 6.45 | Illumina 50 Gb |
| M265-5 | 7B | 79.22% | 2B: 16.88%<br>6B: 3.89% | tall | 6.38 | Illumina 50 Gb |

**Supplementary Table 4.** CDC Stanley pseudomolecule break points for chromosome parts file.

| Chromosome | start | end | Chromosome_part |
|---|---|---|---|
| chr1A | 0 | 295656822 | chr1A_part1 |
| chr1A | 295656822 | 591313643 | chr1A_part2 |
| chr1B | 0 | 352665291 | chr1B_part1 |
| chr1B | 352665291 | 705330581 | chr1B_part2 |
| chr1D | 0 | 247828290 | chr1D_part1 |
| chr1D | 247828290 | 495656580 | chr1D_part2 |
| chr2A | 0 | 401616302 | chr2A_part1 |
| chr2A | 401616302 | 803232604 | chr2A_part2 |
| chr2B | 0 | 395372622 | chr2B_part1 |
| chr2B | 395372622 | 790745243 | chr2B_part2 |
| chr2D | 0 | 328747013 | chr2D_part1 |
| chr2D | 328747013 | 657494025 | chr2D_part2 |
| chr3A | 0 | 379652944 | chr3A_part1 |
| chr3A | 379652944 | 759305888 | chr3A_part2 |
| chr3B | 0 | 428271271 | chr3B_part1 |
| chr3B | 428271271 | 856542542 | chr3B_part2 |
| chr3D | 0 | 314160942 | chr3D_part1 |
| chr3D | 314160942 | 628321883 | chr3D_part2 |
| chr4A | 0 | 377182132 | chr4A_part1 |
| chr4A | 377182132 | 754364263 | chr4A_part2 |
| chr4B | 0 | 348556683 | chr4B_part1 |
| chr4B | 348556683 | 697113365 | chr4B_part2 |
| chr4D | 0 | 252127135 | chr4D_part1 |
| chr4D | 252127135 | 504254270 | chr4D_part2 |
| chr5A | 0 | 357677490 | chr5A_part1 |
| chr5A | 357677490 | 715354979 | chr5A_part2 |
| chr5B | 0 | 356964834 | chr5B_part1 |
| chr5B | 356964834 | 713929667 | chr5B_part2 |
| chr5D | 0 | 286471564 | chr5D_part1 |
| chr5D | 286471564 | 572943128 | chr5D_part2 |
| chr6A | 0 | 313479595 | chr6A_part1 |
| chr6A | 313479595 | 626959190 | chr6A_part2 |
| chr6B | 0 | 357857111 | chr6B_part1 |
| chr6B | 357857111 | 715714221 | chr6B_part2 |
| chr6D | 0 | 241911561 | chr6D_part1 |
| chr6D | 241911561 | 483823121 | chr6D_part2 |
| chr7A | 0 | 371458899 | chr7A_part1 |
| chr7A | 371458899 | 742917797 | chr7A_part2 |
| chr7B | 0 | 374268330 | chr7B_part1 |
| chr7B | 374268330 | 748536659 | chr7B_part2 |
| chr7D | 0 | 321892491 | chr7D_part1 |
| chr7D | 321892491 | 643784981 | chr7D_part2 |
| chrUn | 0 | 266325606 | chrUn |

**Supplementary Table 5.** Heights of homozygous wild type (*Rht-B13a*) or homozygous mutant (*Rht-B13b*) plants from Cadenza0453 for the S240F amino acid mutation in the RNBS-A motif. P-values were calculated using Student's t-test, $p < 0.01$ ** and $p < 0.001$ ***. Internode P-1 corresponds to the internode below the peduncle, with subsequent internodes numbered by their position from the peduncle.

| | Wild type (cm) | Mutant (cm) | p-value |
|---|---|---|---|
| Spike | 8.2 ± 0.3 | 8.1 ± 0.6 | 0.96 |
| Peduncle | 29.1 ± 1.8 | 25.3 ± 1.9 | 0.003 (**) |
| Internode P-1 | 15.6 ± 2 | 9.6 ± 1.4 | <0.001 (***) |
| Internode P-2 | 11.5 ± 2.7 | 6.3 ± 1.5 | <0.001 (***) |
| Internode P-3 | 6.9 ± 1.1 | 5 ± 0.4 | <0.001 (***) |
| Internode P-4 | 1 ± 0.7 | 1.3 ± 1 | 0.57 |
| Total height | 72.3 ± 3.6 | 55.6 ± 3 | <0.001 (***) |

**Supplementary Table 6.** Plant height of $T_1$ segregating progeny from 4 independent transgenic events in wheat cultivar Fielder carrying the *Rht-B13b* allele from Magnif M.

| S No. | Family | Transgene | Height (mm) |
|---|---|---|---|
| 1 | Control-1 | - | 610 |
| 2 | Control-2 | - | 770 |
| 3 | Control-3 | - | 650 |
| 4 | 1-1 | + | 570 |
| 5 | 1-2 | + | 475 |
| 6 | 1-3 | - | 780 |
| 7 | 1-4 | + | stunted |
| 8 | 1-5 | + | 650 |
| 9 | 1-6 | + | 540 |
| 10 | 1-7 | - | 750 |
| 11 | 1-8 | + | 490 |
| 12 | 1-9 | + | 455 |
| 13 | 1-10 | + | 455 |
| 14 | 2-1 | - | 720 |
| 15 | 2-2 | + | stunted |
| 16 | 2-3 | + | 370 |
| 17 | 2-4 | + | stunted |
| 18 | 2-5 | + | stunted |
| 19 | 2-6 | + | stunted |
| 20 | 2-7 | + | 340 |
| 21 | 2-8 | + | stunted |
| 22 | 2-9 | - | 700 |
| 23 | 2-10 | + | 355 |
| 24 | 2-11* | - (0) | 540 |
| 25 | 2-12* | + (2) | 250 |
| 26 | 2-13* | + (2) | 260 |
| 27 | 2-14* | + (2) | 150 |

| | | | |
|---|---|---|---|
| 28 | 2-15* | + (2) | stunted |
| 29 | 6-1* | + (1) | 160 |
| 30 | 6-2* | + (1) | 435 |
| 31 | 6-3 | + | 515 |
| 32 | 6-4* | + (?) | 560 |
| 33 | 6-5 | + | 210 |
| 34 | 6-6 | + | 150 |
| 35 | 6-7 | + | 510 |
| 36 | 6-8 | + | 520 |
| 37 | 6-9* | - (0) | 740 |
| 38 | 6-10 | - | 660 |
| 39 | 7-1 | + | 460 |
| 40 | 7-2 | + | 395 |
| 41 | 7-3 | + | 465 |
| 42 | 7-4 | + | 485 |
| 43 | 7-5 | + | 480 |
| 44 | 7-6 | + | stunted |
| 45 | 7-7 | + | 520 |
| 46 | 7-8 | + | stunted |
| 47 | 7-9 | + | 440 |
| 48 | 7-10 | + | 150 |

*Plants used to identify transgene copy number using Southern hybridisation, transgene copy number indicated in brackets

#### Supplementary Figures

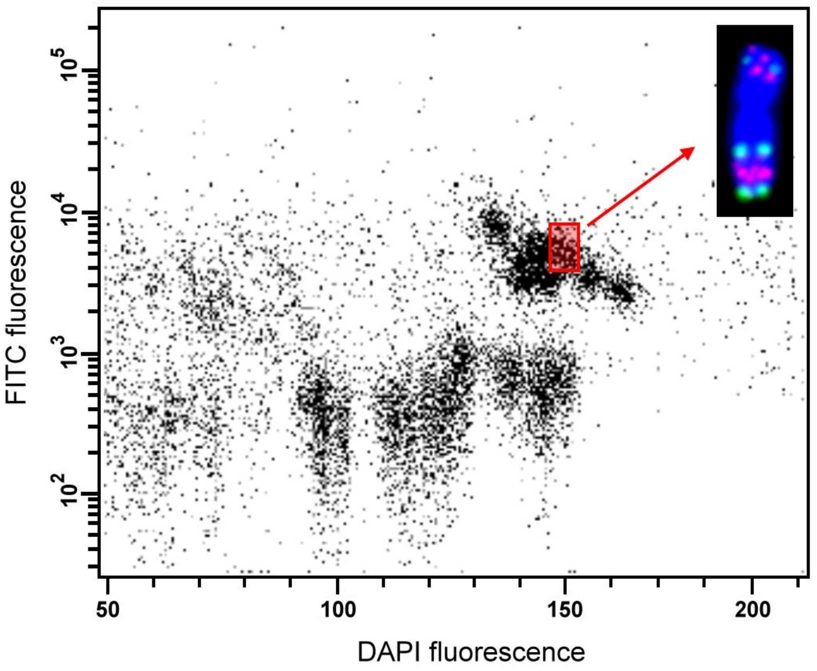


**Supplementary Figure 1**. Bivariate flow karyotype (dot-plot DAPI vs. FITC) obtained after flow cytometric analysis of mitotic metaphase chromosome suspensions prepared from fixed tall and short progeny from a Magnif x Magnif M cross. Prior to analysis, chromosomes in suspension were stained by DAPI and labeled by FISHIS with a FITC-labelled probe for GAA microsatellites. Chromosome 7B carrying the *Rht13* gene was flow-sorted using sort window shown as red rectangle. Inset: Image of chromosome 7B after FISH with probes for pSc119.2 (green) and Afa family (red) repeats; chromosomal DNA was stained by DAPI (blue).

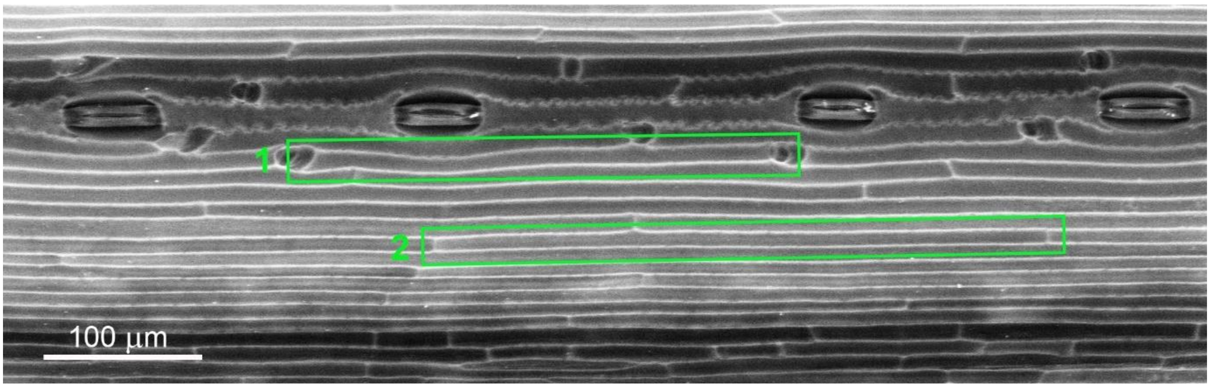


**Supplementary Figure 2**. Scanning electron micrograph of peduncle cells to illustrate cell types measured. 1) inter-hair cell and 2) single cell.

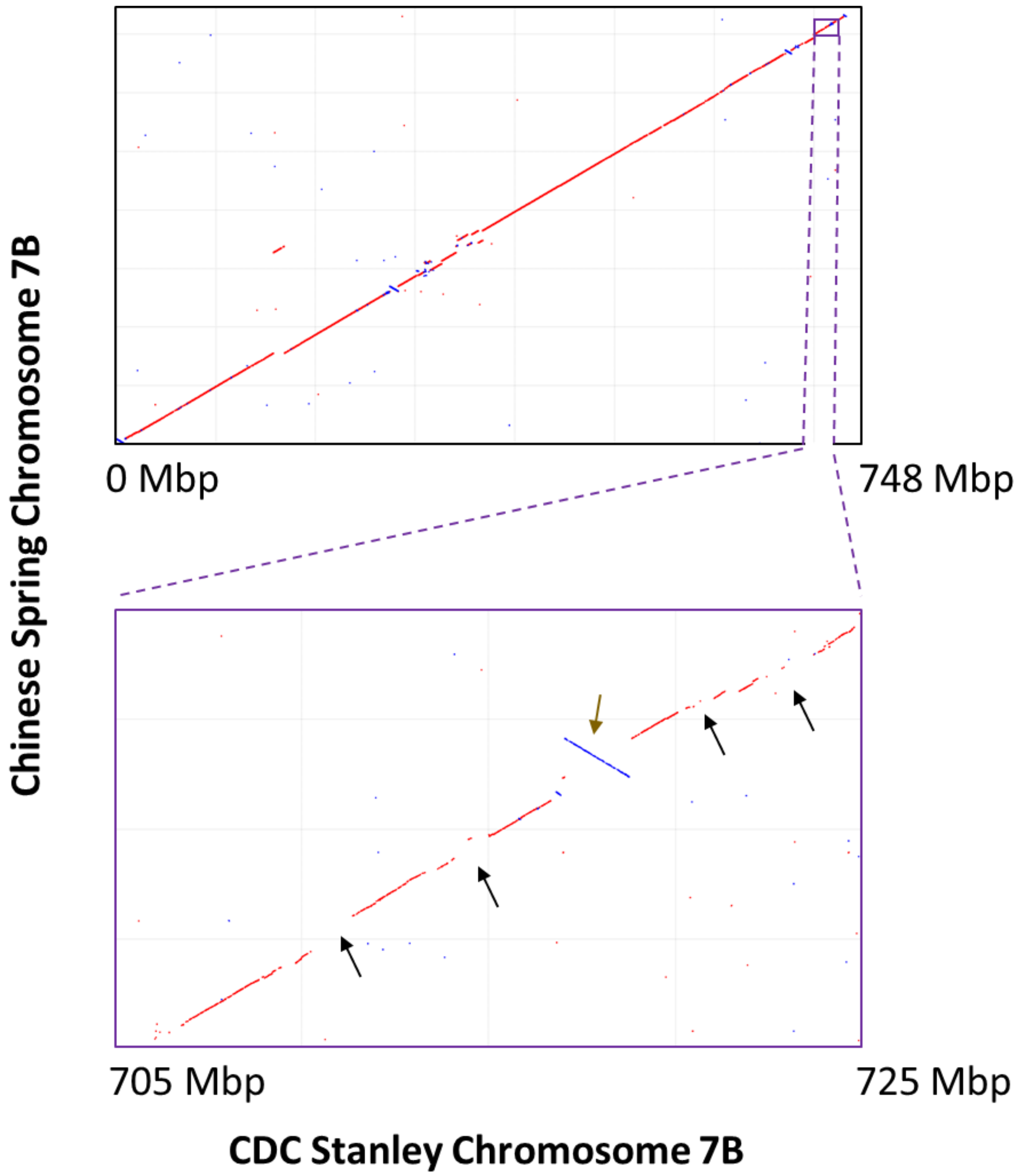


**Supplementary Figure 3.** Dotplot of the alignment between chromosome 7B of CDC Stanley and Chinese Spring. Whole chromosome alignment (top) and localized alignment of the *Rht13* region spanning 705 Mbp to 725 Mbp (bottom) are shown. Putative regions of sequence dissimilarity (black arrows) and an inversion event (gold arrow) are indicated.

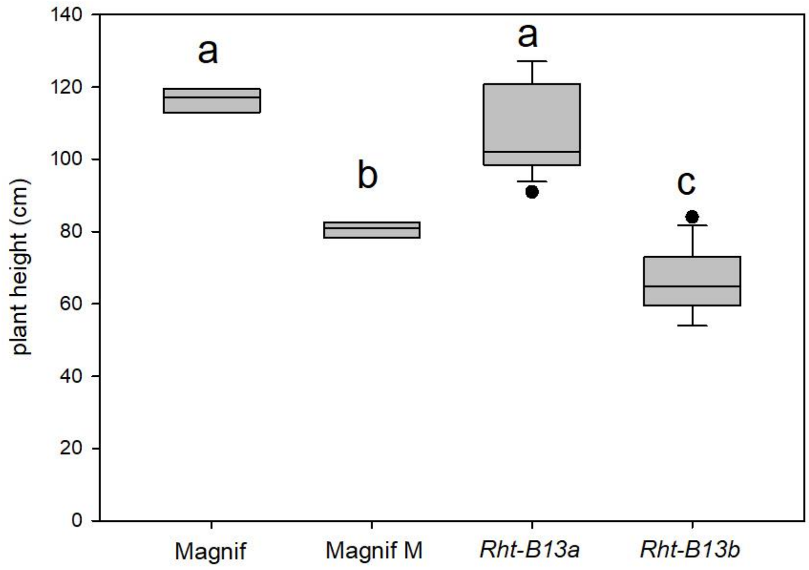


**Supplementary Figure 4.** Plant height of Magnif and Magnif M parental lines and homozygous progeny derived from a cross between Magnif and Magnif M carrying either the *Rht-B13a* or the *Rht-B13b* allele of the KASP marker developed from the SNP found within NB-LRR gene at the *Rht13* locus. Box lower and upper border show the 25th and 75th percentile, line within box shows the median and whiskers correspond to the 10th and 90th percentile. Data points outside the 10th and 90th percentile are indicated as dots. Letters indicate significant differences determined by a one-way ANOVA followed by Tukey post-hoc test (significant level 0.05).

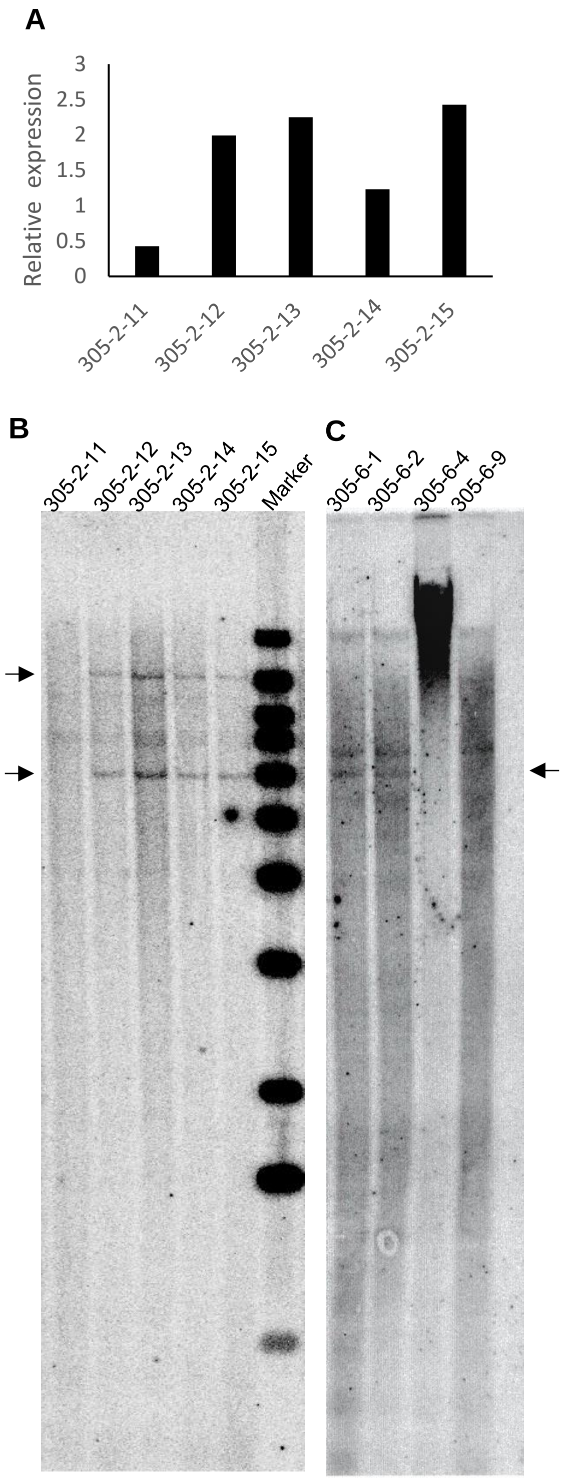


**Supplementary Figure 5.** Analysis of $T_1$ progeny of two transgenic events in Fielder background transformed with *Rht-B13b* allele. (**A**) Relative gene expression of *Rht-B13* in transgenic event #2. (**B**) and (**C**) Southern hybridisation analysis of Fielder transgenics carrying the *Rht-B13b* allele. DNA from $T_1$ segregating lines of transgenic events #2 and #6 were digested with HindIII and a DNA fragment from the selectable marker gene Bar was used as a probe. Arrows indicate the presence of 'Bar' gene in transgenic plants. Copy number is reported in Supplementary Table 5.

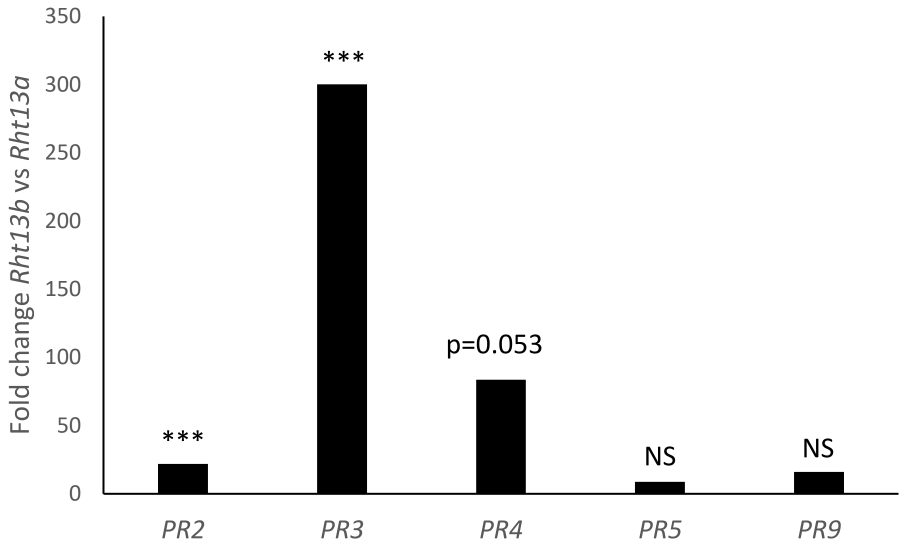


**Supplementary Figure 6**. *PR* gene expression in RNA-seq data from Magnif peduncles from DESeq2 analysis. NS= non-significant, *** $p<0.001$.

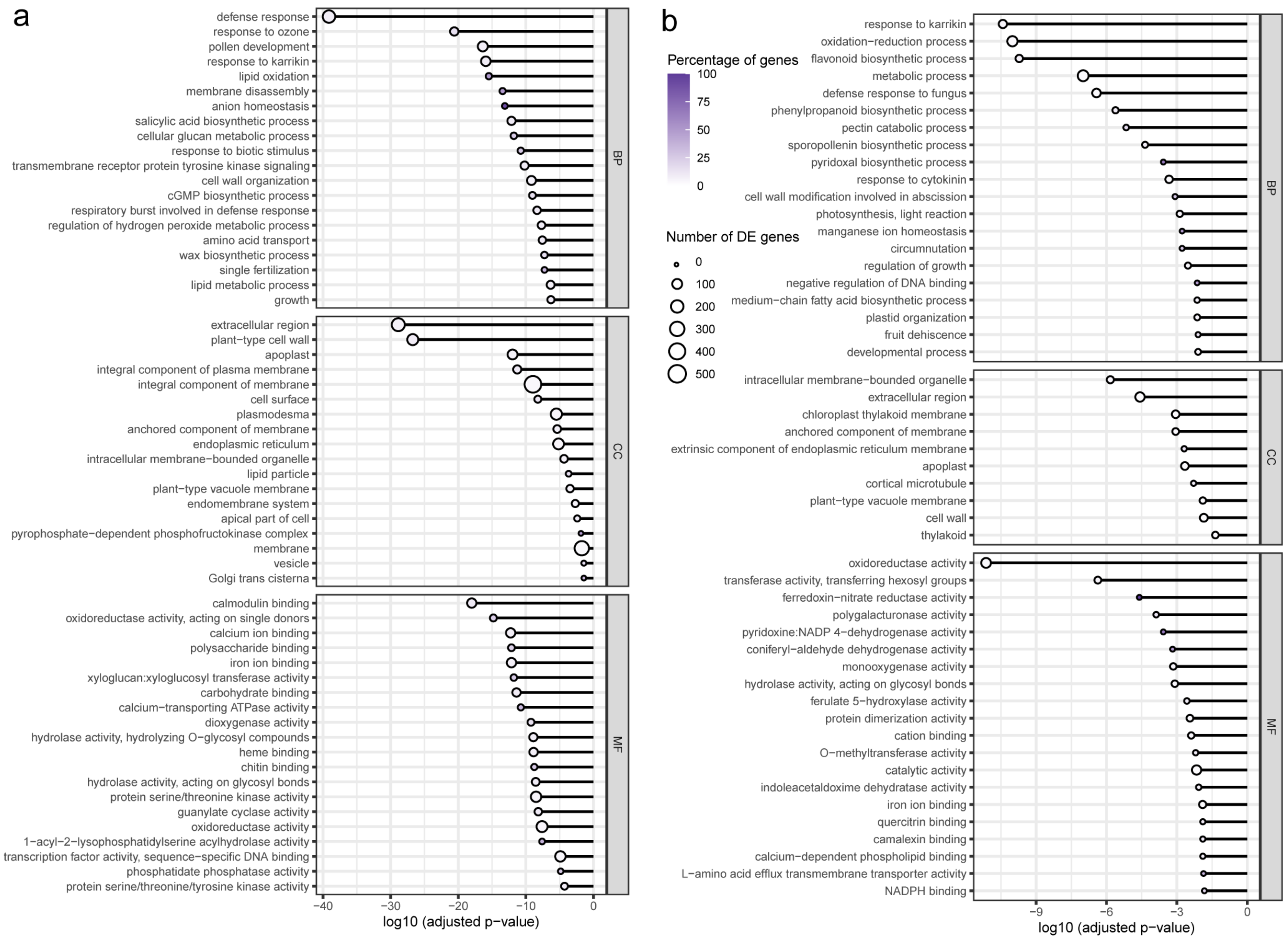


**Supplementary Figure 7.** Analysis of genes differentially expressed in *Rht-B13b* compared to *Rht-B13a* peduncles in a Magnif background. a) GO term enrichment in upregulated genes and b) GO term enrichment in downregulated genes. GO terms were grouped into biological process (BP), cellular component (CC) and molecular function (FF) and Revigo was used to simplify the GO term list, only the top 20 terms by p-value are shown for each category.
